# HENMT1 restricts endogenous retrovirus activity by methylation of 3’-tRNA fragments

**DOI:** 10.1101/2025.05.12.650695

**Authors:** Josh I Steinberg, Helene Sertznig, John J Desmarais, Jenna Wilken, Daisy Rubio, Matthew Peacey, Justin B Kinney, Andrea J Schorn

**Author notes:** These authors contributed equally (Josh I Steinberg, Helene Sertznig).

## Abstract

Long terminal repeat (LTR) retroelements such as endogenous retroviruses (ERVs) utilize host tRNA as a primer for reverse transcription, and are thus susceptible to silencing by small RNAs derived from the 3′-end of mature tRNAs (3′-tRFs). Rigorous quantification reveals that 3’-tRF amounts are not directly proportional to tRNA levels, instead, 3’-tRFs of specific isodecoder tRNAs are highly enriched in a pattern conserved between mouse and human. We found that 3’-tRFs are 2’-O methylated by the small RNA methyltransferase HENMT1 protecting them from degradation and promoting ERV silencing. In the absence of HENMT1, 3’-tRFs are subjected to non-templated tailing by the terminal nucleotidyltransferases TUT4 and TENT2 that regulate small RNA turnover. Due to the perfect sequence complementarity of 3’-tRFs to endogenous retroviral sequences, they have thousands of targets in mammalian genomes. We conducted a massively parallel reporter assay using *Mus musculus particle type D*, a highly active murine ERV, to determine target site rules for 3’-tRFs. Our results suggest that HENMT1 not only stabilizes germline integrity but also serves transposon control by 3’-tRFs in the soma.

## INTRODUCTION

Transposable elements (TEs) are a threat to genome stability. They are usually embedded in inactive chromatin, but epigenetic reprogramming during development and disease erases repressive chromatin marks. This is when small RNA-mediated silencing mechanisms become crucial to regulate TE expression, restrict their mobility and maintain genome integrity. In contrast to miRNAs in animals which function through partial complementarity, small RNAs that detect TEs usually recognize their target by near-perfect sequence complementarity and are stabilized by 2’-O methylation at their 3’-end (Ji & Chen, 2012). This modification is mediated by HENMT1 and its homologs across eukaryotes, and defects in methylation generally lead to destabilization of small RNA substrates (Ji & Chen, 2012; Park *et al*, 2002). In animals, HENMT1 is known for its role in piRNA biogenesis in the germline. Animal miRNA which are not methylated and piRNA that lost 2’-O methylation are prone to untemplated nucleotide addition to their 3’-ends by terminal nucleotidyltransferases, often preceding degradation (Gainetdinov *et al*, 2020; Kamminga *et al*, 2010; Yang *et al*, 2022; Yu & Kim, 2020).

Small RNAs derived from the 3’-end of tRNAs (3′-tRFs) inhibit long terminal repeat (LTR)-retroelements by targeting their highly conserved tRNA primer binding site (PBS) (Li *et al*, 2012; Schorn *et al*, 2017; Yeung *et al*, 2009). The use of host tRNAs as a primer for reverse transcription and replication is a hallmark of LTR-retroelements, which include endogenous retroviruses (ERVs) in mammals but also infectious retroviruses (Chapman *et al*, 1992; Schorn & Martienssen, 2018). In the mouse, ERVs are highly active causing mutations and are expressed during epigenetic reprogramming in stem cells of the developing embryo where also 3’-tRFs levels are high (Gagnier *et al*, 2019; Guo *et al*, 2024; Matsui *et al*, 2010; Rowe *et al*, 2010; Schorn *et al*., 2017). The most active ERV families in mouse, IAP (*Intracisternal A Particle*), ETn (*Early Transposon*), and MusD (*Mus musculus particle type D*), are strongly inhibited by 3’-tRFs (Schorn *et al*., 2017). Different LTR-retroelements have evolved to bind different primer tRNAs and are hence susceptible to 3’-tRFs from that specific tRNA sequence. If the PBS sequence is mutated, the transposon evades silencing, but may lose its ability to replicate. ETn/MusD and HIV-1 replicate using Lysine^3^-TTT or Lys-TTT-3, one of seven Lysine-TTT isodecoder tRNAs that differ only in a few nucleotides.

3’-tRFs are processed from full-length, mature tRNAs including their posttranscriptional CCA-tail and come in two distinct size classes: 22 nucleotides long tRF3b repress coding-competent retroelements post-transcriptionally, while 17-19 nucleotides short tRF3a fragments interfere with reverse transcription (Schorn *et al*., 2017). 3’-tRFs have been found to bind PIWI and AGO proteins in many organisms, both during overexpression and in pulldowns of endogenous proteins (Couvillion *et al*, 2010; Hasler *et al*, 2016; Haussecker *et al*, 2010; Kumar *et al*, 2014; Li *et al*., 2012; Maute *et al*, 2013; Su *et al*, 2022; Yeung *et al*., 2009). Post-transcriptional gene silencing by tRF3b fragments has been demonstrated in reporter gene assays and for a number of genes with target sites in their 3’-UTR (Kuscu *et al*, 2018; Maute *et al*., 2013; Parikh *et al*, 2022; Su *et al*., 2022). The most widespread biological targets of 3’-tRFs are retroviral sequences that have complementarity at the PBS motif in their 5’-UTR (Kawaji *et al*, 2008; Li *et al*., 2012; Schorn *et al*., 2017; Yeung *et al*., 2009), but their target site rules are unknown. In contrast, target site rules of miRNAs and piRNAs have been studied extensively (Bartel, 2009; Gainetdinov *et al*, 2023; McGeary *et al*, 2022; McGeary *et al*, 2019; Vainberg Slutskin *et al*, 2018).

Here, we provide a first systematic analysis of how mutations in the 5’-UTR target site of 3’-tRFs affect the expression of the highly active murine MusD retroelement. Silencing was effective despite mismatches to 3’-tRFs in the region defined as “seed” for miRNAs, but disrupted by mutations outside the seed region reminiscent of piRNAs. Distinct mechanisms regulate small RNA targeting and turnover depending on their sequence complementarity to their targets (Gainetdinov *et al*, 2021; Han & Mendell, 2023). We found 3’-tRFs are stabilized by 2’-O methylation like other small RNA classes with extensive target complementarity. In the absence of 2’-O methylation, 3’-tRFs accumulated adenosine and uridine tails mediated by TENT2 and TUT4, respectively. Terminal RNA modifications of 3’-tRFs and their target site rules suggest that 3’-tRFs are an ancient substrate of the RNAi machinery. We quantified 3’-tRFs and their precursor full-length tRNA molecules and found specific tRFs are highly enriched in a pattern conserved between mouse and human. The availability of full-length tRNA primer and inhibitory 3’-tRFs, their methylation status, as well as mutations at the PBS sequence are constrains that shape the evolutionary success of each ERV family. We found that HENMT1 methylates 3’-tRFs in germ cells but also has a function outside the mammalian germline in repressing LTR-retrotransposition, possibly extending its role to cancer and inflammatory diseases that overexpress TEs.

## RESULTS

### Selective enrichment of 3’-tRNA fragments (3’-tRFs) compared to full-length tRNA

To gain a comprehensive understanding of how tRNA levels relate to 3’-tRF expression in the cell, we isolated mature tRNAs and 3’-tRFs based on size from the same samples profiling different human and mouse stem cell lines (Figure 1A-B). We chose mouse embryonic P19 cells and mouse trophoblast stem cells (mTSC) (Tanaka *et al*, 1998) because of their high expression of ERVs and 3’-tRFs (Maksakova & Mager, 2005; Schorn *et al*., 2017). We also profiled human HeLa cells to observe potential trends across species and because we previously established regulation of the highly active murine ERV families ETn, MusD and IAP by 3’-tRFs in this heterologous host. We quantified full-length, mature tRNAs using hydrolysis-based tRNA sequencing (Gogakos *et al*, 2017) and 3’-tRFs using standard ligation-based small RNA sequencing. Ligation-based small RNA sequencing readily detects 3’-tRFs that carry 5’-phosphate groups akin to miRNA, siRNA, piRNA, and other endonuclease type III cleavage products. Hydrolysis-based sequencing is one of several methods to accurately quantify full-length tRNAs despite their strong secondary structures and abundant RNA modifications that hamper reverse transcription (Gogakos *et al*., 2017; Goodarzi *et al*, 2016; Zheng *et al*, 2015). After optimizing hydrolysis conditions (Extended Data Fig. 1A), hydrolysis products that have 5’-hydroxyl groups are phosphorylated to allow for ligation and cloning. Both, tRNA hydrolysis products and 3’-tRFs, do not readily align to the genome due to their post-transcriptional CCA-tails and require tailored sequence analysis.

**Figure 1.**
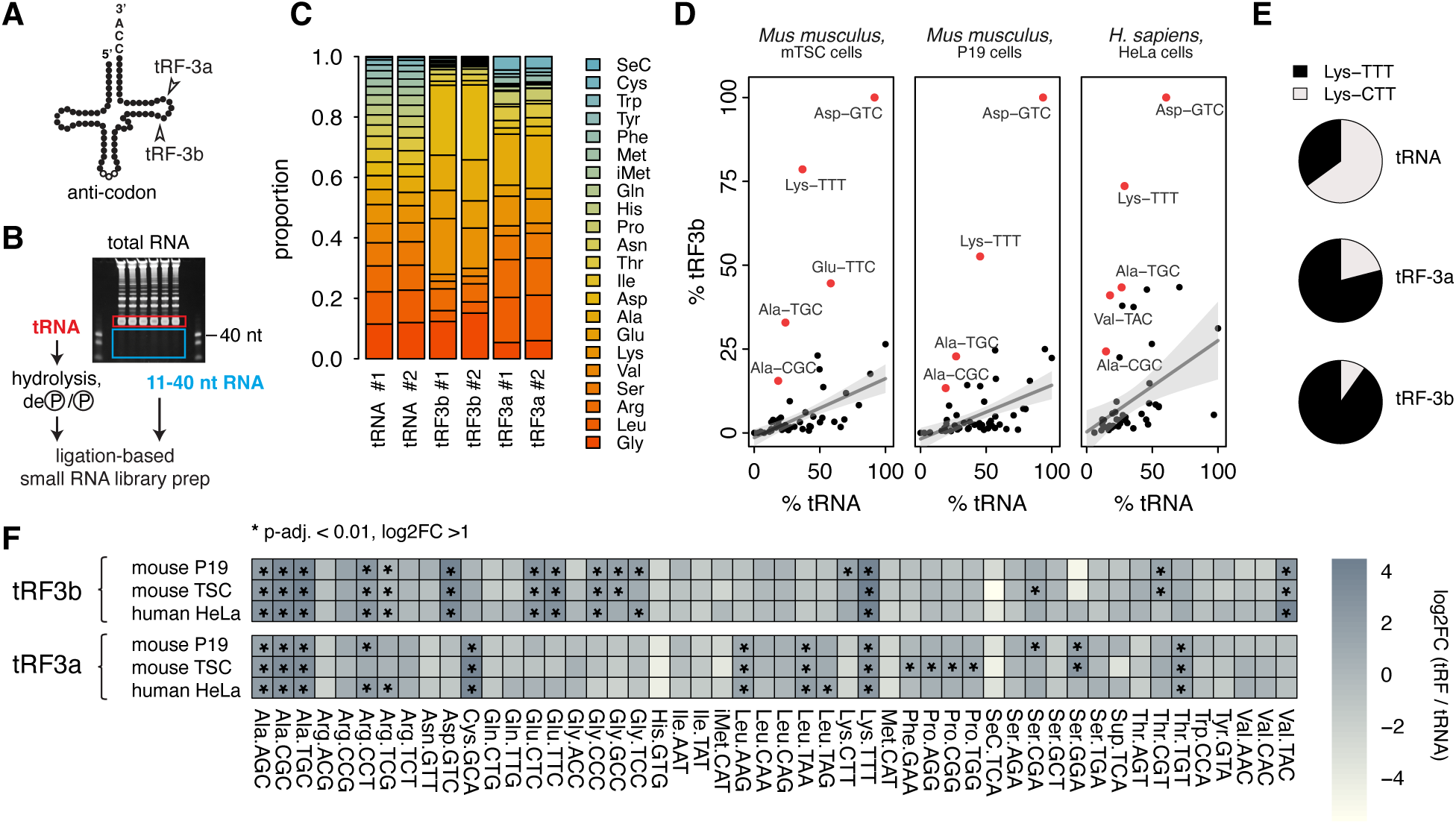
Selective enrichment of 3’-tRFs compared to full-length tRNAs in mouse and human cell lines. **(A)** Diagram of cellular tRF3a (∼18 nt) and tRF3b (22 nt) small RNAs derived from mature tRNAs including their post-transcriptional CCA-tail. **(B)** Work flow of hydrolysis-based tRNA sequencing and small RNA sequencing from the same samples. **(C)** Quantification of full-length tRNAs, tRF3a and tRF3b fragments. Shown are proportions by amino acid isotype in P19 mouse embryonal carcinoma cells, two biological replicates each. **(D)** Relative abundance (%) of tRF3b and full-length tRNA reads ending in CCA. Values are relative to the isoacceptor sequences with the highest read count set to 100%. 3’-tRFs marked in red are highly enriched (log2FC >3, padj. < 0.01) compared to their corresponding full-length tRNA. Enrichment of specific isoacceptors is conserved across mouse and human cell lines. mTSC: mouse trophoblast stem cells. **(E)** Lysine isoacceptor ratios of tRNAs and tRFs in P19 cells. **(F)** Heat map of log2-fold changes (log2FC) of tRF3a and tRF3b relative to their correspond-ing full-length tRNA. Asterisks (*) mark significant enrichment (log2FC >1, padj. < 0.01).

Analysis of tRNA, tRF3a, and tRF3b composition revealed that distinct amino acid isotypes were enriched for each class (Figure 1C, S1B). In general, tRNA composition exhibited a relatively uniform distribution across isotypes with the most highly expressed isotypes representing ∼10-15% of tRNA sequences (Figure 1C, S1B). In contrast, tRFs exhibited a much more skewed distribution, with the most abundant isotypes representing up to ∼26% of 3’-tRFs. Importantly, the composition of 3’-tRFs differed from that of tRNAs and was consistent between replicates (Figure 1C, S1B). Moreover, tRF3a and tRF3b amounts were not related. For example, the most abundant isotype for tRF3a fragments was Alanine across all mouse and human cell lines sampled, while Lysine and Asparagine were the most abundant isotypes for tRF3b. Isotype tRNAs further split up into distinct isoacceptor tRNAs that encode the same amino acid but differ in their anticodon. Mapping reads to isoacceptor sequences resulted in distinct read counts demonstrating that the 3’-terminal sequence of each isoacceptor is unique and sufficient for unambiguous mapping of 3’-tRFs. Isoacceptor ratios differ significantly between tRNAs and tRFs (Figure 1E-F). Lysine has two isoacceptor families, Lys-CTT and Lys-TTT, the latter being the primer tRNA for several retroviruses including HIV-1 and ETn/MusD. While Lys-TTT is slightly underrepresented in the tested cell lines compared to Lysine-CTT, it starkly dominates the pool of tRF3a and tRF3b (Figure 1E).

Remarkably, tRFs of specific isoacceptors were highly enriched in a pattern conserved between mouse and human (Figure 1D, 1F, S1C, S1D). tRF3b fragments of eleven isoacceptors were significantly enriched in all three cell lines and enrichment of eight isoacceptors was shared for tRF3a (Figure 1F). While Alanine (all isoacceptors) and Lysine-TTT enrichment was observed for both tRF3b and tRF3a, other isoacceptor sequences were enriched specifically for either tRF3b or tRF3a. It should be noted that for the majority of isoacceptor sequences, 3’-tRF levels were somewhat proportional to the amount of their parental, full-length tRNA (Figure 1D). In addition to accumulation of 3’-tRFs from specific tRNAs, some tRFs were expressed less than expected. 3’-tRFs significantly underrepresented in all three cell lines for both, tRF3a and tRF3b, were SeC-TCA, Met-CAT, Ile-TAT, His-GTG (Figure 1F). Within each isoacceptor group, different isodecoder tRNAs exist that differ only in a few nucleotides but share the same anticodon. They too exhibit differential enrichment indicating that 3’-tRF amounts are highly regulated (Extended Data Fig. 1E). Lys-TTT-3 or Lys^3^-TTT, the exact isodecoder that primes reverse transcription of HIV-1 and ETn/MusD, was significantly enriched across all cell types for both tRF3a and tRF3b (Extended Data Fig. 1E-F). Given that tRFs from certain tRNA isodecoders that differ only by a few nucleotides are much more abundant than others, this is unlikely due to selective endonucleolytic cleavage of tRFs from tRNAs, but rather suggests selective stability and turnover mechanisms.

### 3’-tRFs are 2’-O methylated and depend on the small RNA methyltransferase HENMT1

We set out to examine post-transcriptional RNA modifications of 3’-tRFs that might account for selective turnover and stability of 3’-tRFs and license them for *bona fide* small RNA silencing. 3’-tRFs have a strong 5’-nucleotide preference for uridine with tRF3b fragments ending in pseudo-uridine of the TψC-arm of tRNAs. Interestingly, pseudo-uridine in small RNAs of the plant germline affects transport and inheritance and also marks select miRNAs and piRNAs in animals (Herridge *et al*, 2025). Small RNAs with extensive sequence complementarity to their targets are stabilized by 2’-O methylation at their 3’-terminus: these include siRNAs in yeast, plants and invertebrates, as well as piRNAs in animals, but not miRNAs in animals that have been adapted for gene regulation and function by partial complementarity. 3’-tRFs have extensive complementarity to their ERV targets owing to the conserved tRNA PBS motif. We analyzed mouse testis small RNA that was oxidized to enrich for 2’-O methylated RNA (Yang *et al*, 2019b) and found strong enrichment of 3’-tRFs (Figure 2A-C) akin to piRNAs that are known to be 2’-O methylated. Of 34 tRF3a and 49 tRF3b isoacceptors expressed above background (>5 BaseMean counts), 13 and 29, respectively, were significantly enriched (log2FC > 1 and padj. < 0.01). For this analysis, we included all sequences annotated as tRNA-derived in the UCSC repeat mask. These are named after the codon, i.e., Lys-AAA loci represent Lys-TTT tRNAs. Both, tRF3a and tRF3b Lysine-TTT fragments, were more than 8-fold enriched, indicating these sequences were robustly methylated. Notably, there is some heterogeneity among piRNA sequences as has been reported by others (Gainetdinov *et al*., 2021). Metazoan miRNAs are generally not 2’-O methylated and were depleted in oxidized samples as expected (Figure 2A). Similar to piRNAs, select 3’-tRF isotypes were heavily affected by oxidation. Ser, Thr, Pro, and Ala tRF3b, as well as Gly and Ser tRF3a fragments, were significantly depleted, thus providing evidence that their 3’-ends were not modified by methylation.

**Figure 2.**
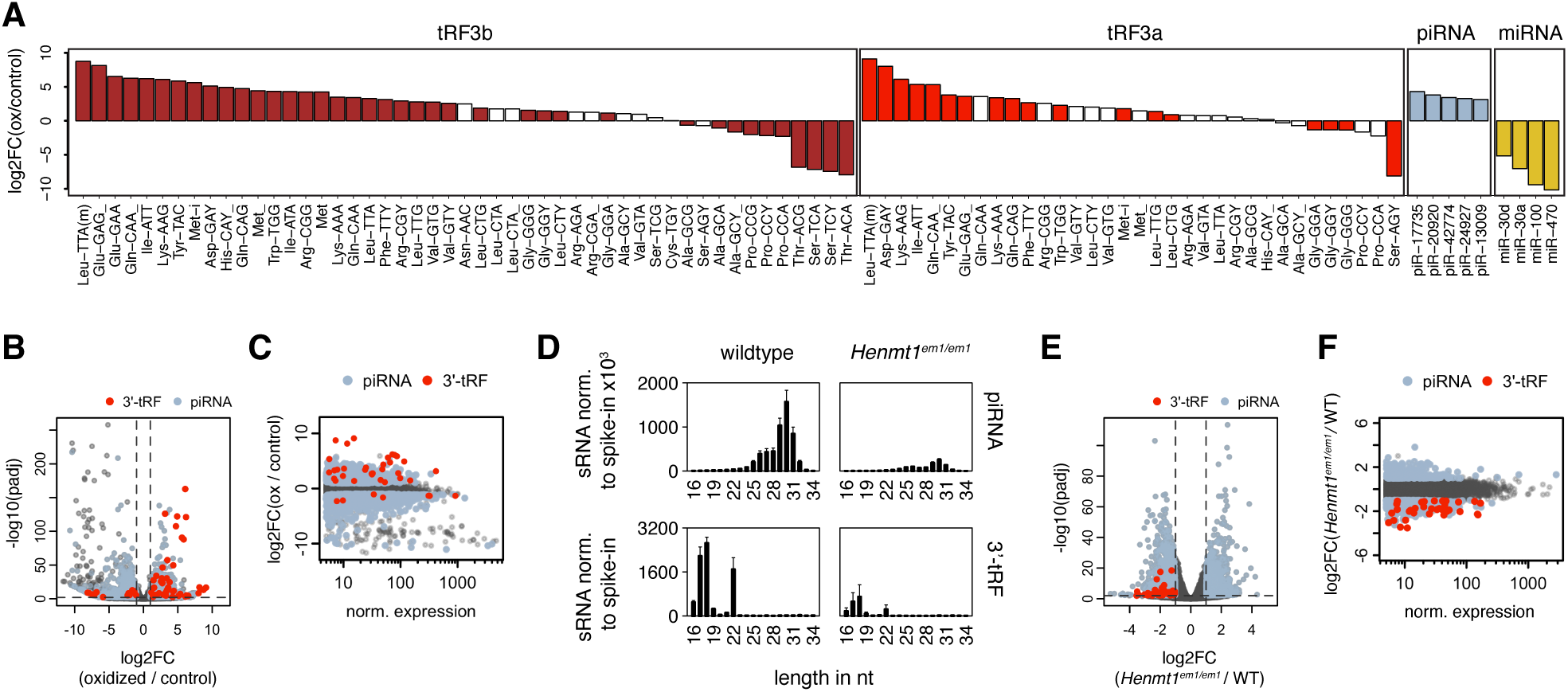
3’-tRFs are stabilized by HENMT1-mediated 2’-O methylation. (**A**) Small RNAs with 2’-O methylation at their 3’-end enrich during periodate oxidation treat-ment. Small RNAs oxidized (ox) versus untreated (control) from mouse testis are shown ranked by their log2-fold change (log2FC). White bars correspond to 3’-tRFs that were expressed but did not exhibit statistically significant changes. For miRNAs and piRNAs, only a few example sequences are shown. (**B**) Volcano plot showing enrichment of 3’-tRFs after oxidation treatment of mouse testis samples. Each dot represents one sequence (piRNA, miRNA) or isoacceptor (3’-tRFs). Signifi-cantly enriched or depleted (Log2FC > |1| and padj. < 0.01) piRNAs and 3’-tRFs are colored in steel-blue and red, respectively. (**C**) Enrichment/depletion of oxidized samples versus control plotted against normalized expression of small RNAs. (**D**) Size distribution of piRNAs (upper panel) and 3’-tRFs (lower panel) in Henmt1^em1Pdz/em1Pdz^ (henceforth abbreviated Henmt1^em1/em1^) and wildtype mouse primary spermatocytes [dataset Gainetdinov et al. 2021]. (**E**) Volcano plot showing depletion of 3’-tRFs in Henmt1^em1/em1^ compared to wildtype primary spermatocytes. Each dot represents one sequence (piRNA, miRNA) or isoacceptor (3’-tRFs). Significantly enriched or depleted (Log2FC > |1| and padj. < 0.01) piRNAs and 3’-tRFs are colored in steel-blue and red, respectively. (**F**) Enrichment/depletion in Henmt1^em1/em1^ versus wildtype (WT) primary spermatocytes plotted against normalized expression of small RNAs.

2’-O methylation of small RNAs is mediated by HENMT1 and is known to support piRNA stability (Horwich *et al*, 2007; Kamminga *et al*., 2010; Kirino & Mourelatos, 2007; Kurth & Mochizuki, 2009; Lim *et al*, 2015; Modepalli *et al*, 2018; Ohara *et al*, 2007; Pastore *et al*, 2021; Saito *et al*, 2007). Indeed, tRF3a and tRF3b were strongly depleted in Henmt1*^em1Pdz/em1Pdz^*loss-of-function mouse germline tissues (Figure 2D-F) (Gainetdinov *et al*., 2021). Small RNA length distribution profiles show loss of piRNA, but also 18 nt tRF3a and 22 nt tRF3b fragments in male Henmt1*^em1Pdz/em1Pdz^* germ cells (Figure 2D, S2C-D. 3’-tRFs were significantly destabilized in primary spermatocytes (spI), secondary spermatocytes (spII), as well as round spermatids (rs) of Henmt1*^em1Pdz/em1Pdz^* mice (Figure 2E, 2F, S2B. Trimming of piRNA precursors during biogenesis is mediated by the 3’-to-5’ exonuclease PNLDC1. While piRNAs are susceptible to degradation in the absence of either HENMT1 or PNLDC1 (Extended Data Fig. 2D) (Gainetdinov *et al*., 2021), 3’-tRFs levels were not significantly affected by PNLDC1. This is consistent with the 3’ ends of 3’-tRFs uniformly consisting of their post-transcriptional CCA-tail.

### In the absence of HENMT1, 3’-tRFs are tailed by terminal nucleotidyltransferases

In the absence of 2’-O methylation, piRNAs accumulate non-templated nucleotide tails at their 3’-ends (Kamminga *et al*., 2010; Lim *et al*., 2015; Svendsen *et al*, 2019). 3’-tails that accompany degradation of miRNAs are mediated by the terminal nucleotidyltransferases TENT2, TUT4 and TUT7 (Han & Mendell, 2023). We analyzed tail length of miRNA, piRNA and 3’-tRFs in male germ cells of *Henmt1^em1Pdz/em1Pdz^*and *Pnldc1^em1Pdz/em1Pdz^* mice (Figure 3A, S3D) (Gainetdinov *et al*., 2021). As expected, neither loss of function had any effect on miRNAs which are not methylated in animals and are not trimmed by PNLDC1 (Figure 3A). In the absence of the trimmer enzyme, piRNAs accumulate a high percentage of 1-5 nt long tails. Tails did not accumulate on piRNAs in *Henmt1^em1Pdz/em1Pdz^*, potentially because trimmer is still there to remove them or because only a fraction of piRNAs are 2’-O methylated and therefore affected. In contrast, 3’-tRFs accumulate tails of 2-3 nt length in either mutant and for both tRF3a and tRF3b fragments. Mono-, di-, and tri-nucleotide additions to 3’-tRFs increased up to three-fold in *Henmt1^em1Pdz/em1Pdz^*germ cells. When looking at the nucleotide composition of tails, miRNA and piRNA tails are dominated by single uridines, while 3’-tRFs carry longer U-tails (Figure 3B). Interestingly, 3’-tRFs in spermatogonia exhibited adenine tailing in addition to untemplated uridine, a pattern also observed for miRNA. Spermatogonia are also the only tissue sampled in which 3’-tRFs were not depleted upon Henmt1 loss (Extended Data Fig. 2B-C), again suggesting 3’-tRFs in this tissue are more miRNA-like.

**Figure 3.**
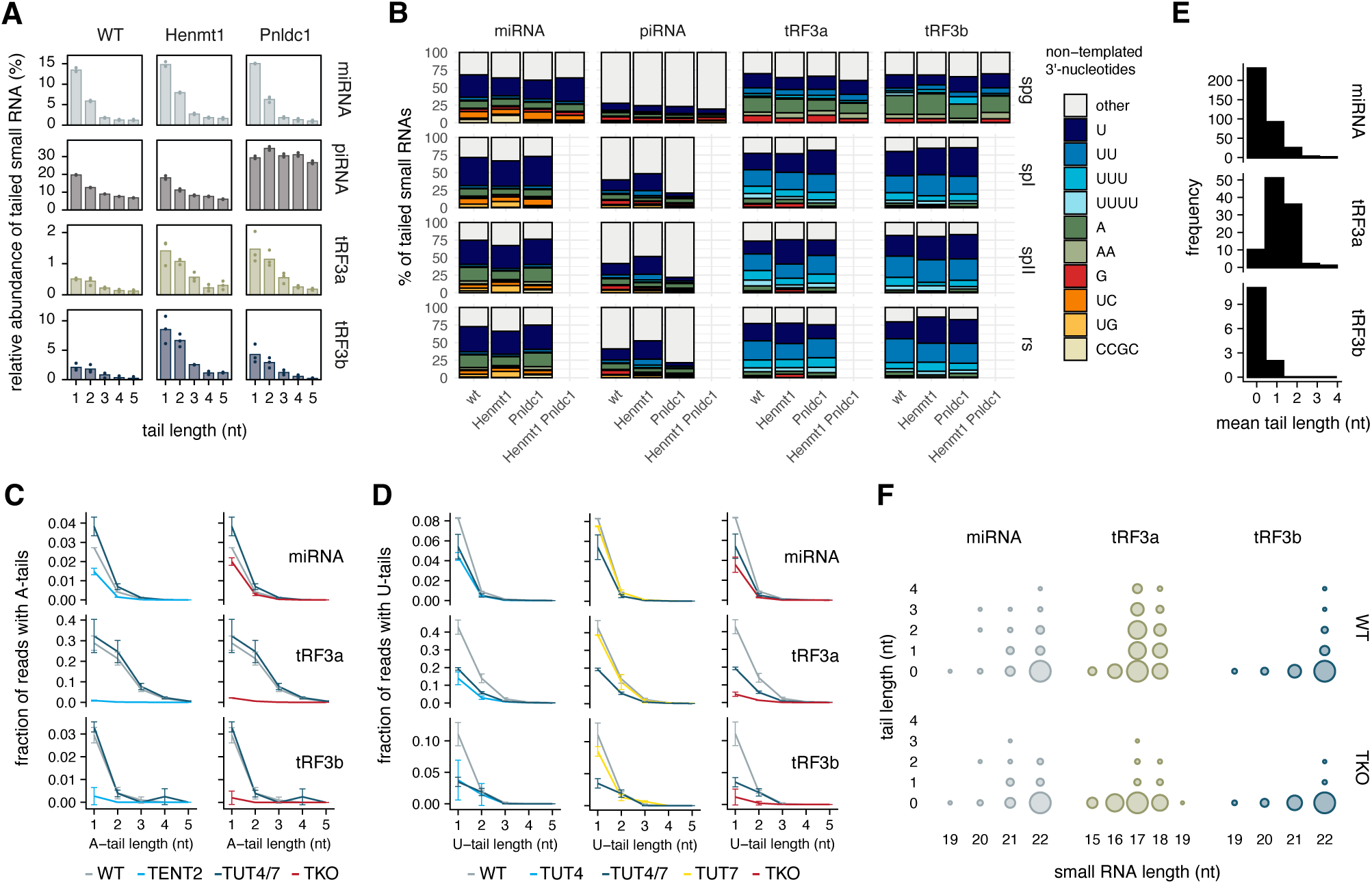
In the absence of HENMT1, 3’-tRFs are tailed by the terminal nucleotidyltrans-ferases TENT2 and TUT4. **(A)** In male germ cells of Henmt1^em1/em1^ mice, 3’-tRFs accumulate tails of non-templated nucleo-tides. Tail lengths are shown for small RNA classes in wildtype (WT) primary spermatocytes and loss of function mutants for Henmt1 and the 3’-exonuclease Plndc1. Relates to Figure S3D. **(B)** Nucleotide composition of untemplated 3’-nucleotides in wildtype and loss of function male mouse germ cells. Spermatogonia (spg), primary spermatocytes (spI), secondary spermato-cytes (spII), round spermatids (rs). **(C)** Adenosine and **(D)** uridine tail length of AGO2-bound small RNAs in HEK293T wildtype (WT) and knockout cells lacking the terminal nucleotidyltransferases TUT4, TUT7, or TENT2. TKO: triple knockout. **(E)** Frequency of untemplated nucleotides by length for AGO2-bound small RNAs in HEK293T cells. Frequency was defined as the number of miRNAs or isodecoder tRF3a/b (greater than 10 reads per million total reads) relative to the number of untailed small RNAs. **(F)** Overall tail length in AGO2-bound wildtype (WT) versus TUT4/TUT7/TENT2 (TKO) HEK-293T cells relative to length of the small RNA without untemplated tail.

To determine which enzymes mediate tailing of 3’-tRFs in the absence 2’-O methylation, we looked for somatic tissues or cell lines that do not express HENMT1. While HEN1 homologs are expressed in both, germline and soma of plants and invertebrates, HENMT1 in animals is known for its role in the germline only. 3’-tRFs are expressed in early development including the germline (our unpublished data), but also in somatic tissues and many cancer cell lines. Interestingly, publicly available GTEx RNA-seq data (Carithers *et al*, 2015) revealed a range of HENMT1 expression across tissues (Extended Data Fig. 3A). On average, cell lines derived from the cervix, testis, and adrenal gland exhibited the highest median expression, whereas those originating from the nervous system, liver, and kidney displayed the lowest expression. We corroborated these findings by Western blot, showing high expression of HENMT1 in HeLa cells (derived from the cervix), but no detectable expression in HEK293T cells (derived from the kidney) (Extended Data Fig. 3B). Thus, we analyzed small RNA data from AGO2-pulldown samples (Yang *et al*., 2022) of single, double, and triple knockouts of TUT4, TUT7 and TENT2 in HEK293T cells for tailing of 3’-tRF. Adenylation and uridylation were the predominant non-templated nucleotides added to 3’-tRFs in HEK293T cells (Figure 3C-D). Notably, tRF3a fragments in wildtype cells exhibited comparable amounts of adenylation and uridylation, whereas both miRNAs and tRF3b showed a two-fold bias towards uridylation. Strikingly, a cumulative of more than 40% of tRF3a reads had non-templated tails at their 3’-end, while only ∼10-14% of tRF3b and miRNAs showed evidence of A- or U-tailing (Figure 3C-D). As previously reported, TENT2 mediates miRNA adenylation and its knockout (KO) resulted in a reduction of A-tails (Figure 3C). The impact on 3’-tRFs was even more pronounced, with a near complete abolition of A-tailing. Uridylation substantially decreased upon the depletion of TUT4 but did not decrease upon loss of TUT7 (Figure 3D). Accordingly, TUT4/7 double-knockouts (DKO) essentially show TUT4 uridylation level. Uridylation of tRF3a fragments was further diminished in the triple knockout (TKO) suggesting that TENT2 can uridylate them in the absence of other enzymes, underscoring its promiscuity for nucleotide incorporation in non-templated tails that has been reported before. Tail length and composition were the same when comparing AGO1- or AGO2-bound small RNAs to total, small RNA (Extended Data Fig. 3E). While the majority of miRNAs and tRF3b fragments were not tailed, tRF3a fragments were on average mono-tailed with up to three nucleotides (Figure 3E-F) and loss of all three transferases resulted in a substantial reduction of tRF3a tails relative to wildtype (Figure 3F). 3’-tRFs from certain tRNA isoacceptors such as Lys-TTT exhibited longer tails than others (Extended Data Fig. 3F-G). In summary, we observed substantial tailing of tRF3a fragments in the absence of HENMT1. Adenine tails of 3’-tRF depend on TENT2 with particularly long A-tails for tRF3a fragments, while uridine tails are short (1 nt) and depend on TUT4.

### HENMT1 has a function outside the animal germline and enforces silencing of LTR-retrotransposition

To determine whether 3’-tRF methylation by HENMT1 has a function in transposon silencing, we used retrotransposition assays. The coding-competent MusD6 co-mobilizes the non-coding ETnIIbeta retrotransposon by recognition of their shared, flanking LTR sequences (Ribet *et al*, 2004). The reporter gene becomes active only after splicing, reverse transcription and integration into the genome (Figure 4A). Retrotransposition assays are performed in a heterologous host to avoid confounding background of thousands of endogenous transposon copies in the original host species. We and others previously established HeLa cells as a heterologous host that is permissive for ETn/MusD retrotransposition and subject to regulation by 3’-tRFs (Ribet *et al*., 2004; Schorn *et al*., 2017). Human HeLa cells were transfected with the murine ERV reporter plasmids (Figure 4A-B), and selected for antibiotic resistance encoded by the reporter gene. Retrotransposition was quantified as the number of resistant cell colonies.

**Figure 4.**
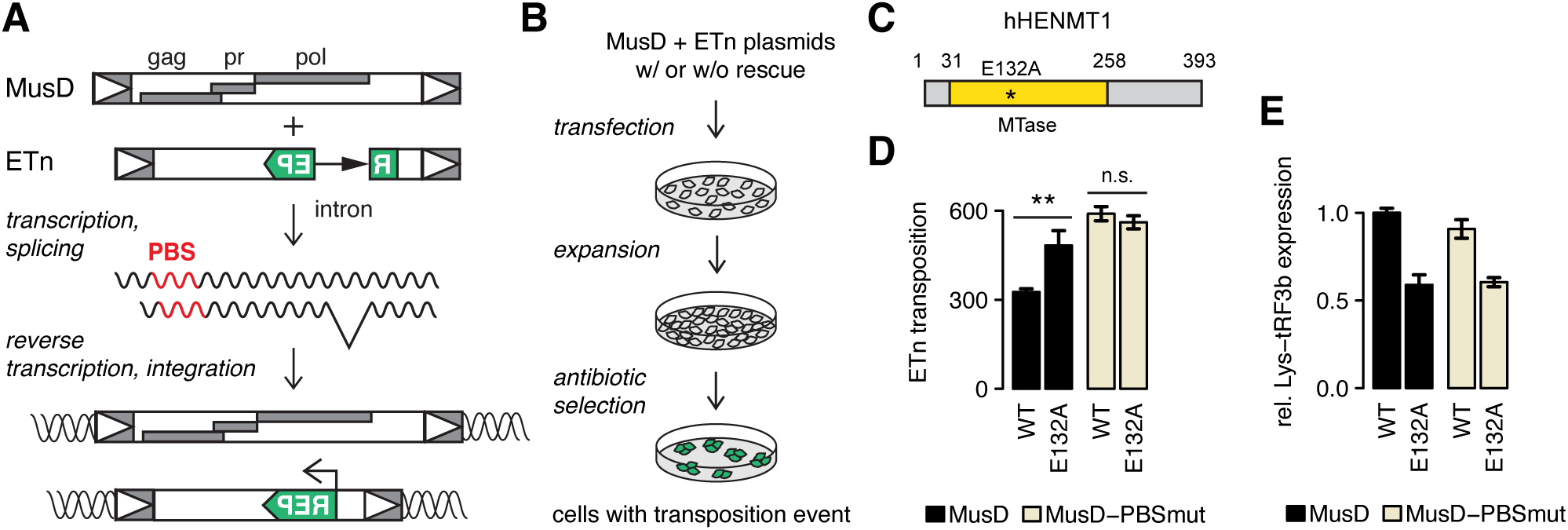
HENMT1 restricts LTR-retrotransposition. **(A)** Plasmid-based retrotransposition assay: The gene products of *Mus musculus particle type D* (MusD) reverse-transcribe the retroviral RNA of MusD and *Early Transposon* (ETn) and integrate their cDNA into the host genome. The reporter (REP) gene becomes active only after splicing, reverse transcription and integration into the genome. Gag: group-specific antigen, pr: protease; pol: polyprotein containing reverse transcriptase/ribonuclease H and integrase domains; PBS: tRNA primer binding site. **(B)** HeLa cells are transfected with plasmids, selected for antibiotic resistance (reporter gene) and retrotransposition is quantified (number of resistant cell colonies). **(C)** Domain structure of human HENMT1 with the E132A mutation indicated, which disrupts methyltransferase (MTase) activity. **(D)** Retrotransposition assay in a clonal Henmt1 CRISPR KO HeLa cell line (clone #29) with expression of functional, wildtype HENMT1 or the catalytic dead HENMT1-E132A mutant, respectively. Loss of HENMT1 methyltransferase activity increased MusD-mediated ETn retro-transposition dependent on the MusD PBS sequence, the target site for 3’-tRFs. Values are mean of colony counts +/- standard deviation. **(E)** Small RNA quantitative PCR with custom primer-probes to detect Lys^3^-tRF3b for samples shown in (D). Relative expression was normalized to miR-191-5p. Values are mean expression +/- standard error.

Indeed, knockdown of HENMT1 resulted in increased retrotransposition (Extended Data Fig. 4A) dependent on the PBS of MusD, the target site for Lysine 3’-tRFs, suggesting HENMT1 controls transposition by methylation of 3’-tRFs. While deletion of the ETn PBS sequence would abrogate tRNA priming for reverse transcription and hence transposition of the ETn-reporter, the MusD PBS sequence can be modified to “escape” regulation by 3’-tRFs with MusD gene products still retrotransposing the ETn-reporter *in trans* (Schorn *et al*., 2017). We tested whether increased transposition is due to increased expression and post-transcriptional regulation of MusD dependent on the PBS in the 5’-UTR using luciferase reporter constructs (Extended Data Fig. 4B). Only the reporter with an intact 18 nt PBS sequence was affected by HENMT1 knockdown. In order to test functional rescue of HENMT1 activity, we generated stable HENMT1 knockout clones (Extended Data Fig. 4C-D). We tested eight CRISPR guide RNAs targeting HENMT1 protein domains for their knockout efficiency by Western Blot (Extended Data Fig. 4C) and generated clonal cell lines (Extended Data Fig. 4D). Henmt1-deficient cell lines show reduced Lys^3^-tRF3b levels (Extended Data Fig. 4E). In order to specifically test the methyltransferase activity of HENMT1, we transiently expressed wildtype and catalytically inactive Henmt1-E132A (Figure 4C) (Peng *et al*, 2018) in Henmt1-deficient cells. The E132A mutation abolishes methyltransferase activity and specifically relieved silencing of transposition dependent on the 3’-tRF target site of MusD (Figure 4D). Lys^3^-tRF3b levels were also rescued in cells with wildtype HENMT1 expression (Figure 4E). We confirmed these results for different amounts and ratios of the coding-competent MusD and the non-coding ETn-reporter (Extended Data Fig. 4F). In all conditions tested, ETn retrotransposition in Henmt1-deficient cells was only affected when driven by wildtype MusD with the intact PBS target site for 3’-tRFs. In contrast, MusD with mutation in the PBS (MusD-PBS^mut^) escapes 3’-tRF regulation and the increased MusD protein amounts drive increased retrotransposition of the ETn reporter (Schorn *et al*., 2017). ETn retrotransposition mediated by MusD-PBS^mut^ was unaffected by the HENMT1 functional rescue (Figure 4D, S4F).

### 3’-tRF target sites in the mouse genome and target site rules for active endogenous retroviruses

Due to the perfect sequence complementarity to endogenous retroviral sequences, 3’-tRFs have many thousands of potential target sites in mammalian genomes. Under stringent criteria allowing no more than three mismatches, tRF3b fragments have ∼9,000 sites of sequence complementarity in the mouse genome (Figure 5A). 5,047 of those are antisense to a repeat annotation (rmsk), 714 are potential target sites in annotated Gencode transcripts, 618 sites are within a Gencode transcript overlapping a repeat annotation, and 2,590 map to non-coding sequences on either strand. Lysine^3^ 3’-tRFs primarily have target sites in TE-derived, repetitive sequences (Figure 5B). Of those 221 transposon-derived target sites, a total of 215 are sequences of the ETn superfamily, which includes MusD6 (ETnERV2-int) and ETnIIbeta (MMETn-int) (Figure 5C). As expected, the large majority of transposon targets are ERVK elements by virtue of their Lysine (K) PBS.

**Figure 5.**
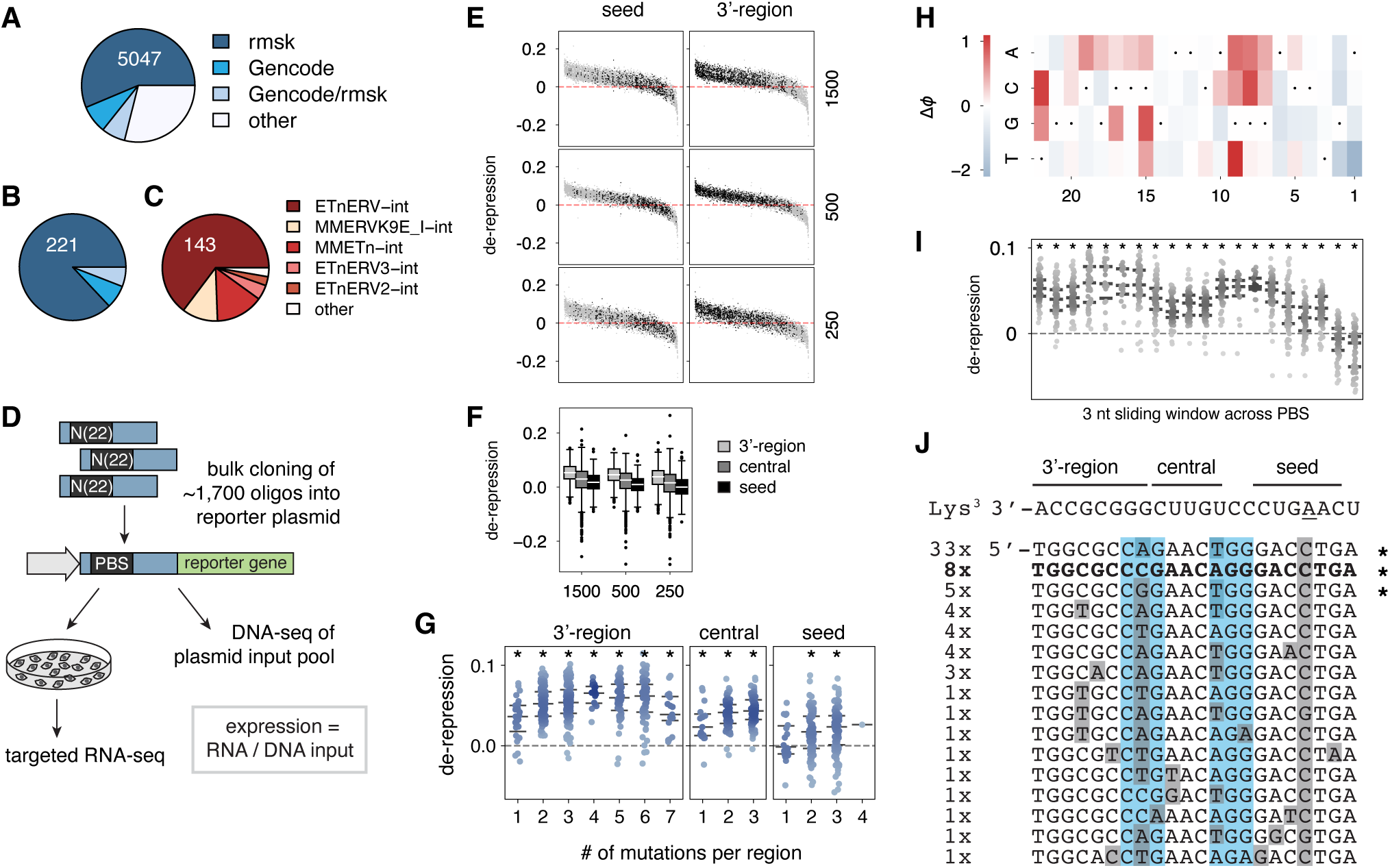
Massively parallel reporter assay (MPRA) to determine 3’-tRF target site rules. **(A)** tRF3b target sequences in the mouse genome (mm10) allowing up to three mismatches. tRF3b target sites within repeats (UCSC rmsk), Gencode transcripts and Gencode transcripts with repeat overlap. “Other” includes tRF3b alignments to either genomic strand. **(B)** Lysine^3^-TTT tRF3b target sequences in the mouse genome categorized as in (A). **(C)** Target sites within repeat sequences in (B) per transposon family. All families except “other” are of the ERV-K (Lysine) superfamily. MusD6 is an ETnERV2-int retroelement and MMETN-int retroelements include ETnIIbeta tested for retrotransposition in this study. **(D)** Experimental scheme of the MPRA to test the effect of ∼1,700 PBS sequences on RNA expression. The MusD6 5’-UTR with the PBS sequence precedes a reporter gene. RNA expression is normalized to DNA abundances in the input plasmid pool. **(E)** Ranked, normalized RNA expression of all biological replicates (three independent plasmid pools transfected at 250, 500 and 1500 ng, respectively). De-repression is relative to the reference canonical PBS sequence (bold sequence in panel J). Seed, central and 3’-region are defined in respect to miRNA nomenclature (depicted in panel J). Each dot (grey) represents one PBS sequence. Black dots indicate PBS sequences with mutations in the seed (left panel) or 3’-region (right panel). **(F)** The same data as in (E) but grouped by whether mutations are in the seed, central, or 3’-region of the PBS. Whiskers represent minimum and maximum datapoints less than 1.5x the interquartile range above or below the quartiles; all outliers are plotted. **(G)** Observed de-repression dependent on the number of mismatches relative to the canonical refer-ence PBS sequence and grouped by region within the PBS. Stars indicate significant de-repression (p-adj. <0.05, 1-sample t-test); lines represent median and quartile values; all data points are plotted. **(H)** MAVE-NN latent phenotype (ΔΦ) map with low (blue) to high (red) expression. Nucleotides of the reference PBS sequence are marked by dots. Positions are aligned with the sequence overview in (J). **(I)** Effect of mutations in the PBS sequence on expression (in 3 nt sliding windows). Positions are aligned with the sequence overview in (J). Lines represent median and quartile values; all data points are plotted. **(J)** The PBS sequences of full-length MusD retroelements in the mouse genome were aligned and ranked by their copy number (indicated on the left). The sequence of Lys^3^-TTT tRF3b is depicted on top of the alignment. The canonical reference PBS sequence (bold) is perfectly complementary to the Lys^3^-TTT sequence except for the conserved mismatch at the m^1^A position (underlined adenosine). Positions with mismatches relative to Lys^3^-TTT are shaded grey. Seed, central and 3’-region were defined as indicated. Nucleotide positions that still allowed replication and priming by Lys^3^-TTT tRNA when mutated in HIV-1 are colored blue. PBS sequences of MusD elements that are known to actively retrotranspose are marked by asterisks.

How many mismatches in the PBS would disrupt post-transcriptional silencing by 3’-tRFs, but still allow for successful tRNA priming and proliferation of the retroelement? We previously analyzed the highly active murine ERV families IAP, MusD, and ETn for sequence conservation at their PBS (Cullen & Schorn, 2020) and noticed that many highly active elements do not have PBS sites that perfectly match their tRNA primer. This suggests successful elements evolve to avoid silencing at the PBS. We wanted to systematically test how complementarity to 3’-tRFs affects expression of ERVs. We collected all MusD full-length elements that fulfilled the following criteria: (i) ERVK:ETnERV-int entries (UCSC Golden Path rmsk annotation) larger than 6 kb, that (ii) are flanked by two LTRs of the ETn superfamily, and that (iii) have a PBS sequence immediately downstream of their 5’-LTR. Twenty-one of those had an intact Gag ORF as well as a PBS site among the top 4 most abundant PBS sequences (Figure 5J). Remarkably, we found that for all these full-length elements, sequence complementarity between the Lys^3^-TTT tRNA primer extends further downstream of the 18 nt PBS up to position 22. This suggests that this ERV family may actually use a 22 nt PBS for replication and certainly is able to base-pair with 22 nt long tRF3b fragments during post-transcriptional silencing. All elements have an A-to-C mismatch at the position that corresponds to the highly conserved 1-methyl-adenosine (m^1^A) position in tRNAs (Figure 5J).

Which mismatches in the PBS would disrupt post-transcriptional silencing by 3’-tRFs? We employed a massively parallel reporter assay (MPRA) (Kinney & McCandlish, 2019) to determine which positions and nucleotides might be critical for 3’-tRF mediated post-transcriptional silencing. We used a reporter plasmid that is sensitive to silencing by 3’-tRFs and contains the 5’-UTR of MusD6 including its PBS upstream of a luciferase reporter gene (Schorn *et al*., 2017). We tested ∼1,700 sequence variations at the Lys^3^-TTT PBS motif. We included the PBS sequences from our analysis of the mouse genome, but also systematically introduced mutations in sliding windows of 3 nucleotides along the extended 22 nt PBS sequence. In addition, we used the miRNA target prediction software miRanda (Enright *et al*, 2003) to design scrambled sequences with poor hybridization scores for binding to Lys^3^-tRF3b. Cells were transfected with bulk-cloned, pooled reporter plasmid DNA, after which RNA was extracted to measure expression using a targeted RNA-seq protocol (Figure 5D). RNA expression was normalized to input DNA amount and quantified by NGS sequencing.

In analogy to nomenclature for base pairing between miRNA guide and target mRNA (McGeary *et al*., 2022), we defined a “seed” region (guide position g2-8), “central region” (g9-13) and “3’-supplemental region” (g14-22) along the 3’-tRF small RNA guide and its PBS target site (Figure 5J). In contrast to miRNAs, mismatches in the seed region did not necessarily relieve silencing (Figure 5E-F). Rather, mutations in the 3’-region de-repressed MusD-reporter expression. De-repression was defined as an increase compared to the “canonical” PBS sequence, which is perfectly complementary to its Lys^3^-tRNA primer except for the conserved mismatch at the m^1^A position. RNA expression levels were highly reproducible between biological and technical replicates (n = 9) (Figure 5E, S5A-C). A few sequences were over- or under-represented in sequencing read depth, i.e., abundance (Extended Data Fig. 5B), among them the parental plasmid with the MusD6 wildtype 5’-UTR, but overall library skew is low across all replicates and conditions (Extended Data Fig. 5C). Library skew was quantified using a Lorenz curve which displays the evenness in the distribution of sequencing reads across library members, a ll replicates and conditions had a Gini index of less than 0.21 indicating a very even distribution of reads (Extended Data Fig. 5C). In order to correlate RNA to protein expression changes, we cloned a few select reporter plasmids out of the pool and quantified protein expression using the luciferase reporter gene. Of note, the relatively small, but highly reproducible fold changes of RNA expression quantified by the MPRA translated into ∼2-fold changes of protein expression in luciferase assays (Extended Data Fig. 5D). RNA and protein expression for a PBS sequence with zero mismatches to Lys^3^-tRF3b had significantly lower expression than the wildtype MusD6 PBS with three mismatches suggesting that active retroelement copies have evolved to escape silencing. In other words, genomic coding-competent ERV copies with mismatches in their PBS seem to be at an advantage for expression.

To characterize the effects of PBS mutations in more detail, we first grouped sequence variants by the number of mutations per region relative to the canonical reference PBS (Figure 5G). Mutations in the 3’- and central regions strongly increased expression. Increasing mismatches in the central region means a greater central bulge, i.e., unpaired nucleotides between small RNA and target, and further increased expression. Mismatches within the seed region had variable effects on expression. Single mismatches did not significantly change de-repression. Double and triple mutations produced highly variable effects on de-repression but increased de-repression on average, suggesting repression or de-repression depends on nucleotide identity and position. In all regions, expression increased with the number of mismatches between PBS and Lys^3^-tRF3b (Figure 5G, S5E).

Position and nucleotide identity can be scored for their impact on expression using algorithms specifically developed for MPRA. We used the MAVE-NN algorithm to create a genotype to phenotype map (Tareen *et al*, 2022) that scores additive contributions from each possible base at each position to a latent phenotype (Δϕ) that is nonlinearly related to measured expression values (Figure 5H). Changes in ϕ result from mutations disrupting the canonical PBS sequence. Strikingly, nucleotide changes at certain positions had a stark effect on expression. Changes at the target positions t7-9 and t15-22 generally disrupted silencing with increased expression indicated in red. Those positions coincide with highly conserved positions in 3’-tRFs such as the post-transcriptional CCA-tail. Three nucleotide sliding windows along the PBS sequence show de-repression in several areas that are often mutated in genomic MusD copies (Figure 5I). Mutations in the 5’-end of the PBS, close to the 3’-end and CCA-tail of the tRNA primer, will disrupt tRNA priming and replication. Therefore, while advantageous for expression, these elements will not be “successful” and not increase copy number in the genome. However, positions that do not restrict mobility of the retroelement and still allow reverse transcription, can accumulate mutations and be of advantage for high expression. Mutations that still allow tRNA priming are known from actively transposing, high copy number MusD elements (asterisks, Figure 5J) (Ribet *et al*., 2004) and can potentially be inferred (blue shaded) from mutagenesis studies on HIV-1 that uses the same Lys^3^-tRNA primer (Wang *et al*, 2018). PBS mutations that preserve tRNA priming but evade 3′-tRF silencing offer a selective advantage and these elements should accumulate to higher copy numbers in the genome. Indeed, many of the positions mutated in active full-length MusD copies in the mouse genome coincide with positions that conferred de-repression in the MPRA (Figure 5H-J). This suggests that, with many layers of regulation controlling transposition *in vivo*, systematic mutagenesis in an expression-only context as in this MPRA can effectively dissect regulatory constraints on ERVs.

## DISCUSSION

Our data demonstrates a role for HENMT1 in “licensing” 3’-tRFs for transposon control. We found that 3’-tRFs are marked and stabilized by 2’-O methylation (Figure 2) much like other small RNA classes that exhibit extensive sequence complementarity to transposons and viruses, and that serve host defense. The highly active murine ERV MusD is specifically restricted by HENMT1’s methyltransferase activity in retrotransposition assays (Figure 4). Our data indicate that transposon control by 3’-tRFs shares key characteristics with other small RNA silencing pathways. 3’-tRFs are processed from their precursor molecules, full-length tRNAs, in a highly regulated fashion (Figure 1). They share sequences and RNA modifications typical of *bona fide* regulatory small RNAs. This includes their 5’-uridine or pseudo-uridine as well as 2’-O methylation at their 3’-ends. Their transposon targets show signatures of selective pressure to escape 3’-tRF mediated silencing (Figure 5). In the absence of 2’-O methylation, 3’-tRFs become substrates of terminal nucleotidyltransferases (Figure 3) and accumulate A- and U-tails akin to piRNAs and miRNAs (Gainetdinov *et al*., 2020; Kamminga *et al*., 2010; Lim *et al*., 2015; Yang *et al*., 2022).

3’-tRFs showed distinct patterns of enrichment when compared to the composition of parental, full-length tRNAs in the cell (Figure 1), and are clearly not by-products of tRNA decay. In mouse and human, tRNA transcript levels from specific loci vary between cell types whereas tRNA anticodon pools do not, supporting stable translational outcomes (Gao *et al*, 2024; Schmitt *et al*, 2014). Consistent with this, we saw largely similar tRNA expression between human and mouse cell lines when grouped by anticodon (Extended Data Fig. 1C). 3’-tRF production can be manipulated by overexpression of single tRNAs (Kuscu *et al*., 2018), and the abundance of many 3’-tRFs is roughly proportional to their tRNA level (Figure 1C). However, our data clearly shows strong enrichment of specific 3’-tRFs isoacceptor and isodecoder sequences in a pattern conserved between mouse and human (Figure 1D, 1F). Although tRF3a and tRF3b fragments of the Lys^3^-TTT (Lys-TTT-3) isodecoder are significantly enriched compared to their full-length tRNA in all cell lines tested, their expression levels varied starkly between cell lines (Extended Data Fig. 1F). Processive reverse transcription *in vivo* also requires interactions with the tRNA beyond the PBS sequence, and hence constrains LTR-retroelements to use specific isodecoder tRNA sequences that their proteins evolved to bind to (Keeney *et al*, 1995; Le Grice, 2003; Liu *et al*, 2010). Lys^3^-TTT tRNA is expressed at low levels in both mouse stem cell lines, and so may restrict retrotransposition in these cells, despite their high expression level of ETn/MusD (Maksakova & Mager, 2005; Schorn *et al*., 2017; Xu & Boeke, 1990). HeLa cells on the other hand express large amounts of this primer tRNA (Extended Data Fig. 1F) and evidently support reverse transcription of ETn/MusD. Small RNA and 3’-tRF regulation however are permissive to mismatches (Figure 5) and Lys^3^-TTT primed retroviruses may well be regulated by 3’-tRFs of other isodecoder sequences. tRNA *versus* 3’-tRF levels could determine which cell types are permissive to mobility versus expression of the elements. In fact, 3’-tRFs have the capacity to affect both, expression level and reverse transcription in these cell lines. The 3’-end of related isodecoder tRNAs and therefore 3’-tRFs are similar enough in sequence (Extended Data Fig. 1G-H) that one ERV family could be regulated by several isodecoder 3’-tRFs.

tRF3b fragments mediate post-transcriptional silencing (Haussecker *et al*., 2010; Kuscu *et al*., 2018; Maute *et al*., 2013; Schorn *et al*., 2017) and have a large number of predicted target sites in metazoan genomes due to the abundance of LTR-retroelement derived sequences (Figure 5A-B). PBS sequences of active mouse ERV families show signatures of 3’-tRF mediated regulation (Cullen & Schorn, 2020). Mutations in the PBS of mobile copies can be interpreted as selective pressure to escape 3’-tRF mediated silencing (Figure 5J). Our results from the MPRA clearly show that mutations confer an advantage for expression of MusD. Interestingly, there is not a single perfect match of Lys^3^-tRF3b in the entire mouse genome. The most frequently observed mismatch corresponds to the m^1^A in tRF3b fragments, at position 58 of tRNAs. This modification is an A-form RNA helix disrupter (Zhou *et al*, 2016) and might therefore not matter for base-pairing with tRF3b fragments. The observed cytosine in the ETn/MusD PBS is most likely due to the common misincorporation of nucleotides during reverse transcription of m^1^A at this position of the tRNA (Ryvkin *et al*, 2013). It is worth keeping in mind that most TEs in mammalian genomes are non-autonomous. For retroelements that have no “cis” preference to mobilize their own RNA, protein-coding “master-copies” that accumulate mutations at their PBS may lose the ability to prime reverse transcription of themselves but can achieve high expression by evading post-transcriptional silencing and retrotranspose a large number of non-coding, non-autonomous elements with perfect PBS sites. Interestingly, a large number of PBS sequences in inactive and immobile elements are very well conserved. Even though their PBS motif is no longer required for tRNA priming, it may serve co-opted regulation of these transcripts by 3’-tRFs. Many of these elements have essential functions in early development and participate in gene-regulatory networks in mouse and human (Faulkner *et al*, 2009; Franke *et al*, 2017; Guo *et al*., 2024; Modzelewski *et al*, 2021; Peaston *et al*, 2004; Rebollo *et al*, 2012).

Target rules for miRNAs are well established and rules for piRNAs are emerging. In fact, a number of MPRA designs have been used to measure miRNA-target interactions (McGeary *et al*., 2019; Vainberg Slutskin *et al*., 2018). Strong binding between guide (positions g7-15 and g8-16) and targets have been observed for PIWI complexes even in the absence of seed base pairing (Anzelon *et al*, 2021; Gainetdinov *et al*., 2023). This is thought to benefit the ability of PIWI proteins to silence transposons that mutate frequently. Our data on de-repression of the MusD transposon sequence much resemble piRNA target interactions. MusD repression is sensitive to mutations in the 3’-region, but less sensitive to mutations in the seed sequence that dominates miRNA-target interactions (Figure 5). Although the MPRA will include potential transcriptional effects on reporter plasmid DNA by e.g., zinc finger (ZNF) proteins, only a few of the positions with strong effects in the MAVE-NN analysis (Figure 5H) overlap the binding motif of human ZNF417/587, which has evolved as another layer of defense against Lysine^3^-primed retroviruses (Turelli *et al*, 2020). The biggest effects on expression were observed for mutations in the region that base-pairs with the highly conserved CCA-end of 3’-tRFs. Few examples of LTR-retroelements being able to switch between isodecoder tRNAs for priming at low frequency have been observed (Cullen & Schorn, 2020; Das *et al*, 1997; Fennessey *et al*, 2019; Markopoulos *et al*, 2016). Therefore, 3’-tRF target sites are certainly under different evolutionary constraints than miRNA target sites. Also, every round of replication bears a 50:50 chance for *Retroviridae* like MusD of reverting their PBS sequence to a cDNA copy of their primer tRNA (Berwin & Barklis, 1993; Cullen & Schorn, 2020; Hu & Hughes, 2012), further constraining escape from silencing.

3’-tRFs are subjected to A-tailing by TENT2 and U-tailing by TUT4 similar to miRNAs. TUT4 and TUT7 were not redundant (Figure 3C-D), but U-tails on 3’-tRFs were TUT4-dependent. In the absence of 2’-O methylation by HENMT1, extent and length of tailing of tRF3b fragments are similar to that of miRNAs (Figure 3A, 3C-D). Both small RNA classes act in post-transcriptional silencing. We observed long, TENT2-dependent A-tails for tRF3a fragments in HEK293T cells (Figure 3C, 3E-F). A-tailed Lys^3^-tRF3a are perfectly complementary beyond the ETn/MusD target site, and although overall levels are low, it is conceivable that tailed 3’-tRFs regulate an additional target repertoire similar to reports for tailed miRNA (Yang *et al*, 2019a). Untailed tRF3a are rather similar in size to tinyRNAs (tyRNAs) that have been described to guide AGO3-mediated silencing (Sim *et al*, 2023). Similar to overall tRF enrichment patterns, tailing was highly specific to some isodecoders but not to others (Extended Data Fig. 3F-G).

We found that 3’-tRFs are 2’-O methylated and depend on HENMT1 for stability. This is reminiscent of small RNAs in other organisms which depend on 2’-O methylation for stability (Ji & Chen, 2012; Kamminga *et al*., 2010; Kurth & Mochizuki, 2009; Modepalli *et al*., 2018; Zhao *et al*, 2012) and target highly complementary sequences such as transposons and viruses. Phylogenetic analyses show that vertebrates have lost the antiviral AGO clade (Wynant *et al*, 2017). Long, double-stranded RNA (dsRNA) generally invokes an interferon response in vertebrates restricting the generation of small RNAs from dsRNA of repetitive elements. Few tissues such as oocytes in mice and rats produce endogenous siRNAs using a specific Dicer isoform that efficiently cleaves long dsRNA and circumvents RNA immunity pathways (Flemr *et al*, 2013; Tam *et al*, 2008; Watanabe *et al*, 2008; Yang *et al*., 2019b). Oocytes of other mammals express a fourth PIWI protein (PIWIL3) to produce 19 nt short piRNAs (Loubalova *et al*, 2021; Lv *et al*, 2023; Wang *et al*, 2023; Yang *et al*., 2019b). Neither of those oocyte small RNA classes are 2’-O methylated and neither are metazoan miRNAs that evolved to silence targets via partial complementarity. It makes sense that 3’-tRFs are 2’-O methylated, akin to other small RNAs that silence RNA with near perfect pattern match. 3’-tRF might represent an ancient defense mechanism to detect mobile repetitive RNA in early-stage embryonic cell types that cannot be surveilled by the vertebrate immune system.

Our data suggests a functional role of HENMT1 outside the germline of animals. HENMT1 methyltransferase activity restricts ETn/MusD retrotransposition in somatic cells (Figure 4D). Both, HENMT1 and 3’-tRFs are expressed in early development, but also in somatic tissues and in many cancer cell lines. Depletion of HENMT1 results in growth defects and scores as strongly selective across cancer cell lines (Extended Data Fig. 3C). HENMT1 was the top dysregulated RNA-modifying enzyme in a cancer study comparing tumors to matched normal samples (Begik *et al*, 2020). The small RNA methyltransferase is also a prognostic marker in cervical cancer (CESC) according to the TCGA Protein Atlas data (https://www.proteinatlas.org/ ENSG00000162639-HENMT1/cancer). None of these observations can be explained by HENMT1’s role in piRNA biogenesis in the germline. Rather, HENMT1’s role in the soma might be to methylate 3’-tRFs and promote ERV silencing. Somatic ERV re-activation leads to sustained inflammation and is a common theme in aging, cancer and auto-immune diseases (Jansz & Faulkner, 2021; Liu *et al*, 2023; Volkman & Stetson, 2014).

## DATA & CODE AVAILABILITY

Sequencing data have been deposited at *National Center for Biotechnology Information Sequence Read Archive* and are publicly available as of the date of publication using accession numbers GSE296629, GSE296630, and are publicly available as of the date of publication. Original western blotting images have been deposited at *Mendeley* (10.17632/phccbh57wx.1) and are publicly available as of the date of publication. All published software required to reanalyze the data reported in this paper is described in the material and methods section. Code for analyzing MPRA data and producing associated figures can be found at *GitHub* https://github.com/jackdesmarais/TRf_transposon_silencing_MPRA; the release associated with this submission is archived on *Zenodo* at https://zenodo.org/records/15361369 (DOI: 10.5281/zenodo.15361369). Any additional information required to reanalyze the data reported in this paper is available from the lead contact upon request.

## ACKNOWLEDGMENTS

We thank the Cold Spring Harbor Laboratory (CSHL) Core Facilities Genome Targeting and NGS Sequencing, which were supported by the US National Institutes of Health grant P30CA045508. We thank the Chris Vakoc (CSHL, NY) and Janet Rossant (The Hospital for Sick Children, Toronto) laboratories for plasmids and cell lines. We thank Andalus Ayaz from the Kinney laboratory (CSHL) for technical assistance. This work was supported by the US National Institutes of Health (NIH) grant R01 GM138669 to A.J.S. Computational work on the CSHL high performance compute cluster was performed with assistance from the US National Institutes of Health grant S10OD028632-01. J.I.S. was supported by NIHGM MSTP Training award T32-GM008444. D.R. was supported by grant NSF REU-DBI-1950621 to the Undergraduate Research Program (URP CSHL). J.J.D. and J.B.K. were supported by NIH grants GM133777 and HG011787.

## AUTHOR CONTRIBUTIONS

J.I.S. and A.J.S designed the study. J.I.S., H.S., J.W., D.R. and M.P. performed the experiments. J.I.S., J.J.D., H.S., J.W. and A.J.S. analyzed the data and/or its significance. A.J.S. wrote the manuscript with contributions from H.S., J.J.D., J.K.; J.K. and A.J.S. acquired funding.

## DECLARATION OF INTERESTS

The authors declare no competing interests.

## EXTENDED DATA

Extended Data Figures 1-5

Supplemental Table S1

## MATERIAL AND METHODS

### Cell Lines and Tissue Culture

Mouse trophoblast stem cells (TS6.5 derived from B5/EGFP transgenic mice, kind gift from the Janet Rossant laboratory, The Hospital for Sick Children, Toronto, Canada) were grown in MEF-conditioned medium supplemented with FGF4 (R&D systems, 235-F4-025) as previously described (Tanaka *et al*., 1998). P19 mouse embryonic teratocarcinoma cells (ATCC, #CRL-1825) were grown in *Alpha-MEM* (Thermo Fisher Scientific, #12571089) with 7.5% bovine calf serum and 2.5% fetal bovine serum (FBS). HeLa cells (ATCC, #CCL-2) were grown in *DMEM* with 10% FBS. All cell lines were grown at 37C and 5% CO_2_.

### Hydrolysis-based tRNA-sequencing and Small RNA sequencing

RNA was extracted using *TRIzol* (Thermo Fisher Scientific, #15596026) following the manufacturer’s instructions but with 80% v/v ethanol washes during precipitation. 20 ug total RNA per sample were separated on a 6% *Novex TBE-Urea Gel* (Thermo Fisher Scientific, #EC68655BOX) and stained with *SYBR Gold* (Thermo Fisher Scientific, #S11494) to excise tRNAs (visible band ∼72 nt) and small RNAs (11-40 nt, using oligonucleotide markers). RNAs were eluted in 0.3 M NaCl, 0.5 mM EDTA overnight at 4C, ethanol-precipitated in the presence of *GlycoBlue* (Thermo Fisher Scientific, # AM9516), resuspended in a small volume of water and stored in *DNA LoBind Tubes* (Eppendorf, #022431021). The tRNA fraction was treated as previously described (Gogakos *et al*., 2017). Briefly, 250 ng of size-selected tRNA in 125 ul water was combined with 125 ul of 20 mM carbonate-bicarbonate buffer pH 9.8 (Na_2_CO_3_/NaHCO_3_), and RNA was hydrolyzed for 45 min at 90C. After ethanol-precipitation, RNA was treated with *Calf Intestinal Phosphatase* (NEB, #M0290S) for 1h at 37C to remove phosphate groups at the 3’-end of hydrolysis products. The reaction was stopped by acid phenol-chloroform (Thermo Fisher Scientific, #AM9720) extraction, followed by ethanol-precipitation and several washes with ice-cold 80% v/v ethanol to remove excess salt. After resuspension in RNase-free water, RNA was phosphorylated using *T4 Polynucleotide Kinase* (NEB, #M0201S) and cleaned up with acid phenol-chloroform followed by ethanol precipitation before it was used as input in a small RNA library preparation. Small RNA library preparation was performed on both, the treated tRNA fraction and the small RNA fraction of the same samples, using 3’ and 5’ (N)_4_-randomized adapters and PEG-8000 according to *Supplementary Protocol 4* of Giraldez et al. 2018 (Giraldez *et al*, 2018).

### Analysis of hydrolysis-based tRNA-sequencing and 3’-tRFs

Reads were quality filtered using Gordon Assaf’s *fastx-toolkit*. *Cutadapt* (Martin, 2011) was used to clip adapter sequences and size select for 14-60 nt read length. Reads that end in the post-transcriptional CCA-trinucleotide tail of mature tRNA were used for analysis of both, (i) small RNA data, specifically tRF3a and tRF3b fragments ending in CCA, as well as (ii) hydrolysis-based tRNA data, since these CCA-ending reads directly correspond to the amount of full-length parental tRNA of a given isoacceptor or isodecoder. The UCSC *GtRNAdb* annotation (https://gtrnadb.ucsc.edu/genomes/eukaryota/) for mouse (Mmusc10) and human (Hsapi38), respectively, includes isodecoder identity. The post-transcriptional CCA-tail was added to the 3’-end of the mature tRNA sequences in fasta format and used to build a *Bowtie* index (Langmead *et al*, 2009). Reads ending in CCA were directly aligned to this index allowing up to two mismatches to account for RNA modifications (bowtie -f --best --tryhard -v 2). tRF3a fragments were defined as 17-19 nt in length and tRF3b fragments were required to be 22 nt long. Differential expression between 3’-tRF and hydrolysis-based tRNA read counts was calculated using *DESeq2* (Love *et al*, 2014). The pheatmap R package (https://github.com/raivokolde/pheatmap) was used to visualize fold changes of 3’-tRFs relative to their parental tRNA sequences with unsupervised clustering and without clustering, respectively.

Analysis of Lysine isodecoder tRNA sequences (Extended Data Fig. 1F-G) was performed using the same UCSC *GtRNAdb* mature tRNA sequence annotation. Lysine isodecoder tRNA sequences for mouse and human, respectively, were aligned in *Jalview* (version 2.11.4.1) using the *Clustal-Omega* algorithm and default parameters. The *Clustal-Omega* alignment and its distance matrix was used to make a phylogenetic tree in *Jalview* using the neighbor joining method.

### Analysis of small RNA from male mouse germ cells and total testis

Mouse testis small RNAs treated with NaIO_4_ were analyzed for enrichment compared to untreated samples (GSE95218) (Yang *et al*., 2019b). Sorted, male germ cells from wildtype, *Henmt1^em1Pdz/em1Pdz^*, and *Pnldc1^em1Pdz/em1Pdz^* (also referred to as *Henmt1^em1/em1^*and *Pnldc1^em1/em1^*) (PRJNA660633) (Gainetdinov *et al*., 2021) were analyzed for depletion of small RNA classes. Analysis of tRNA-fragments was done as described in Schorn et al. 2017 (Schorn *et al*., 2017). Briefly, reads were split into sequences that end in CCA and reads that don’t. Reads that end in CCA were clipped for their terminal CCA and aligned to the UCSC mm10 genome with everything masked that is not a tRNA in the UCSC rmsk annotation (ftp://hgdownload.soe.ucsc.edu/goldenPath/mm10/database/rmskOutCurrent.txt.gz). *Bowtie2* was used to randomly distribute multi-mapping reads (bowtie2 -N 1 -L 15 --gbar 20 -R10) (Langmead & Salzberg, 2012). Using Bamtools (Barnett *et al*, 2011), sequences were filtered for 0-2 mismatches to allow for RNA modifications common in tRFs. For reads that matched these criteria, tRNA codon, genomic coordinates and read counts were collected, asking for an original read size of 17-19nt (tRF3a) and 20-23 nt (tRF3b). For downstream sequence and size distribution analysis, CCA-tails were added back after the alignment. To analyze tRNA reads that are not 3’-tRFs, reads that did not end in CCA were mapped to the genome with everything except tRNA sequences masked, as described above. tRFs that span splice-sites are not captured by this approach. Again, tRNA codon, loci and read counts were collected for downstream analysis.

Reads that did not map tRNA loci were collected for further analysis of other small RNA classes and intersection with repeats. Reads that aligned to a hairpin precursor (miRBase version 22) (Kozomara *et al*, 2019) with 1 mismatch were classified as miRNAs. Reads that did not match any previous category, were aligned to the UCSC mm10 mouse genome using *Bowtie2* (bowtie2 -N 1 -L 15 --gbar 20 -p2 -f -R10) distributing multi-mappers randomly, and intersected with the current USCS *rmsk* repeat annotation using *BEDtools* to determine repeat- and transposon-derived small RNA (Langmead & Salzberg, 2012; Quinlan & Hall, 2010). Subsequently, reads were intersected with the piRNA annotation piRNAdb.mmu.v1.7.6 (https://www.pirnadb.org) (Rosenkranz *et al*, 2022) asking for at least >90% overlap. To avoid over-counting when collapsing on piRNA names (several previously annotated piRNAs can overlap one read), *BEDtools grouby* was used (bedtools groupby -grp 1-4 -c 5 -o collapse).

The following raw read counts were collected and used for *DESeq2* analysis to determine significantly enriched or depleted small RNAs: 3’-tRFs, other tRFs (sequences that do not end in CCA but map to genomic tRNAs), miRNAs and piRNAs with or without repeat-overlap. To obtain size distributions, reads normalized to spike-in calibrators per number of cells were plotted by length.

### Analysis of untemplated, 3’-end nucleotide tails of small RNAs

Analysis of 3’-tailing of small RNAs utilized data from male germ cells of wildtype, *Henmt1^em1Pdz/em1Pdz^*, and *Pnldc1^em1Pdz/em1Pdz^*mice (PRJNA660633) (Gainetdinov *et al*., 2021), as well as of HEK293T cells with loss of function mutations in the terminal nucleotidyltransferase genes TUT4, TUT7, and TENT2 (GSE183384) (Yang *et al*., 2022). Small RNAs were mapped to an index of *miRBase22* mature miRNAs (Kozomara *et al*., 2019) and the last 22 nt of mature tRNA sequences including their CCA-tail (generated from UCSC *GtRNAdb* as described above). The m^1^A RNA modification in 3’-tRFs often leads to misincorporation of nucleotides during the reverse transcription step of sequencing protocols. To allow alignment to the index without mismatches, this position was replaced by degenerate nucleotides (A/G/T/C). To analyze non-templated tailing of small RNAs, an iterative approach was used: initially, all reads were mapped to the index and those that aligned with zero mismatches were counted as total, untailed small RNA. Reads that did not align, including those with a tail, were shortened by exactly one nucleotide at their 3’ end, remapped, and counted as a tail length of +1 nt if they aligned with no mismatches in this iteration. This process was repeated seven times, each time recording the nucleotide that was clipped from the 3’ end to keep track of tail composition. I.e., tail length reflects the number of iterations that were required to map a read.

### Western Blots

Cells were lysed in ice-cold RIPA buffer: 150 mM NaCl, 1% v/v NP-40, 0.1% w/v SDS, 0.5% w/v sodium deoxycholate, 50 mM Tris-Cl pH 8, 1x Complete mini protease inhibitor (Roche, #11836170001). DTT (50 mM) and bromophenol blue (0.013% w/v) were added to the lysates before heat denaturation. Proteins were separated by SDS-PAGE and transferred onto a 0.45 um nitrocellulose membrane (Biorad, #1620115). Membranes were blocked in 5% w/v non-fat dry milk (Santa Cruz Biotechnology) in 1x TTBS (150 mM NaCl, 20 mM Tris-Cl pH 7.5, 0.05% v/v Tween-20). Human HENMT1 was detected using primary antibody (Thermo Fisher, #PA5-55866, polyclonal rabbit, antigen sequence: MEENNLQCSS VVDGNFEEVP RETAIQFKPP LYRQRYQFVK NLVDQHEPKK VADLGCGDTS LLRLLKVNPC IELLVGV, corresponding to N terminal amino acids 1-77 of human HENMT1) diluted 1:250 in TTBS with milk overnight at 4C and secondary HRP-coupled antibody (Jackson Immuno Research Lab, #111-035-144). Images were developed using Pierce ECL Western Blotting Substrate (Thermo Fisher Scientific, #32106) and the Odyssey Fc imaging system (Li-Cor). Membranes were stripped with Restore Western Blot Stripping Buffer (Thermo Fisher Scientific, #21059) and reprobed for GAPDH or ATCB, respectively (Actin beta, Rabbit, Abcam, #ab8227; GAPDH, Mouse, Abcam, #ab8245).

### CRISPR-knockout and rescue of HENMT1

The puromycin resistance gene in pLenti-Cas9-P2A-puro (Addgene #110837) was replaced by the zeocin resistance gene of pcDNA3.1(+)-zeo. Lentiviral particles carrying Cas9-zeo were harvested from HEK293T cells (ATCC #CRL-3216) after calcium phosphate transfection of pLenti-Cas9-P2A-zeo, pCMV-VSV-G (Addgene #8454), psPAX2 (Addgene #12260 Didier Trono lab). HeLa target cells were infected at low MOI and selected for zeocin to produce Cas9-expressing stock cells. CRISPR guide RNAs (gRNAs) against *HENMT1* were designed using the CSHL domain-focused algorithm (https://mojica.cshl.edu/) and cloned into LR2.1T-puro (kind gift of Chris Vakoc lab, CSHL), a derivative of LRG2.1-Puro (Addgene Plasmid #125594). All gRNA sequences are listed in Supplemental Table S1. HeLa-Cas9 target cells were infected with lentivirus carrying gRNAs to induce targeted mutations and tested for HENMT1 expression by Western Blot. To generate stable clonal cell lines, HeLa cells were infected at low MOI to express hHENMT1_gRNA_#2 (Supplemental Table S1) and selected for puromycin resistance. ∼40 colonies were hand-picked to expand clonal cell lines for cryo-preservation and to collect protein lysate for HENMT1 Western detection. Genomic DNA was extracted from candidate clones (Invitrogen PureLink Genomic DNA Mini Kit, #K182001) and CRISPR-induced mutations were confirmed by Sanger sequencing of PCR-amplified genomic DNA (primers hHenmt1_gDNA_g2_#1/#2) of ∼300 bp around predicted gRNA target sites. Sanger sequencing traces of multiple alleles were resolved (Fig.S4D) using the ’CRISP-ID’ algorithm (Dehairs *et al*., 2016).

*HENMT1* cDNA (NM001102592) was purchased from *Transomics* and cloned into the pcDNA3.1(+)-neo backbone (Thermo Fisher Scientific, #V79020) for transient expression. The cDNA was mutated to be resistant to CRISPR gRNA cleavage, but without changing the amino acid sequence of HENMT1. Plasmid sequences are listed in Supplemental Table S1. The catalytic point mutation (E132A) (Peng *et al*., 2018) was introduced by primer-mediated PCR mutagenesis using high fidelity polymerases (Phusion High-Fidelity DNA Polymerase, NEB #M0530L and KOD Xtreme Hot Start DNA Polymerase Sigma-Aldrich, #71975-M) and subsequent digest with DpnI (NEB, # R0176S). All DNA amplified by PCR or assembled using *NEBuilder HiFi* (NEB #E5520) was confirmed by Sanger-sequencing or *Plasmidsaurus* high coverage NGS sequencing service. Varying amounts of *HENMT1* expressing plasmids were transfected and proteins were checked by Western Blot for their expected molecular weight. For functional rescue of HENMT1, 900 ng of pcDNA3.1(+)-neo-*HENMT1* plasmids per 12-well were co-transfected with transposon plasmids.

The HENMT1 expression data in human tissues used for the analysis in this manuscript (Extended Data Fig. 3A) were obtained from the GTEx Portal on 04/09/2025, v10 release. Gene dependency data (Extended Data Fig. 3C) were obtained from the DepMap Public 24Q1 release (https://depmap.org) (Tsherniak *et al*, 2017), using CRISPR-Cas9 gene effect scores from the Achilles project (Avana dataset). Dependent cell lines were defined as those with a gene effect score < –0.5 for the gene of interest, as per DepMap criteria.

### Transposition assays

The neomycin resistance cassette in pCMV-ETnIIbeta3-TNFneo (Ribet *et al*., 2004) that contained a mirtron was replaced by a SV40-driven blasticidin resistance gene with Tk-poly(A) interrupted by a human globin intron (M91037.1) in reverse orientation (amplified from pJJ101-L1.3-blast, kind gift of the John Moran laboratory, University of Michigan). For each transposition assay, 1.5 x 10^5^ HeLa cells were seeded in triplicates in 12-well plates the day before transfection. 20 ng pCMV-MusD6 (Ribet *et al*., 2004) and 200 ng pCMV-ETnIIbeta3-blast plasmids per 12-well were co-transfected using *Lipofectamine2000* (Thermo Fisher Scientific, #11668019). Transposon sequence accession numbers are AC124426 and AC126548, respectively. ETnIIbeta3 is most closely related to the MMETn-int consensus sequence of the *RepeatMasker* annotation; MusD6 is an ETnERV2-int element. 1-2 days post transfection, cells were transferred onto 10 cm plates and put under blasticidin selection starting day 5-6 until complete. Blasticidin-resistant colonies were fixed with 10% v/v formaldehyde in PBS, stained with 0.1% w/v methylene blue in PBS, and counted (each colony = one retrotransposition event).

### siRNA-mediated knockdown of HENMT1

For HENMT1 knockdown experiments, *ON-TARGETplus Human HENMT1 siRNA SMART pool* (Dharmacon, # L-016309-02) were co-transfected with the above-described transposon plasmids to a final concentration of 25 nM. A 1:1 mixture of *ON-TARGETplus Non-targeting Control siRNAs* #2 (Dharmacon, #D-001810-02) and #3 (Dharmacon, #D-001810-03) was used as negative control. Oligonucleotide sequences are listed in Supplemental Table S1.

### Quantitative PCR RNA analysis

RNA was extracted using *TRIzol* (Thermo Fisher Scientific, #15596026) following the manufacturer’s instructions but with 80% v/v ethanol washes during precipitation. Total RNA was treated with *TURBO DNaseI* (Thermo Fisher Scientific, #AM1907). Reactions were stopped by heat inactivation after addition of EDTA and cleaned up using *AMPure XP beads* (Beckman Coulter, #A63880). For long RNA quantification, RNA was reverse transcribed using *SuperscriptIII* (Thermo Fisher Scientific #18080044) following the manufacturer’s instructions and quantified using *PowerTrack SYBR Green Master Mix* (Thermo Fisher Scientific, #A46109). Reactions were run on a *QuantStudio 6 Flex Real-Time PCR System* (Thermo Fisher Scientific). For small RNA quantification, reverse transcription was performed using *TaqMan Advanced miRNA cDNA Synthesis Kit* (Thermo Fisher Scientific, #A28007) and *TaqMan Fast Advanced Master Mix* (Thermo Fisher Scientific, #4444556), following the manufacturer’s protocol. A total of 10 ng of purified RNA was used for cDNA synthesis, which included polyadenylation, adapter ligation, reverse transcription, and *miR-Amp* pre-amplification steps. Quantitative PCR was carried out on a *QuantStudio 6 Flex Real-Time PCR System* (Thermo Fisher Scientific) using a custom-designed *TaqMan Advanced miRNA Assay* (Thermo Fisher Scientific, #A25576) specific for Lys^3^-TTT tRF3b (5’-UCAAGUCCCUGUUCGGGCGCCA-3’). C_t_ values were normalized to the endogenous control miRNA, miR-191-5p, using the ΔΔC_t_ method. All primer and probe sequences are listed in Supplemental Table S1.

### Genome-wide 3’-tRF target site analysis

Mature tRNA sequences of the mm10 mouse genome annotation were downloaded from the UCSC table browser (’Genes and Gene Predictions’/’tRNA genes’). After adding a CCA-trinucleotide for the post-transcriptional 3’-tail, the last 22 nt were used to generate a fasta file of tRF3b fragments. Sequences were aligned against the UCSC mm10 genome allowing up to three mismatches (bowtie -f --all -v 3) (Langmead *et al*., 2009). Aligned sequences were intersected with transcript (Gencode version M25) or repeat annotations (ftp://hgdownload.soe.ucsc.edu/goldenPath/mm10/database/rmskOutCurrent.txt.gz) using BEDTools (intersectBed -wb -f 1.0) (Quinlan & Hall, 2010). The sequence of the 22 nt tRF3b fragment derived from the Lysine^3^-TTT isodecoder is 5’-UCAAGUCCCUGUUCGGGCGCCA-3’ and its target sequences are the reverse complement. In other words, target sites in transcripts or repeat-derived RNA are defined by tRF3b alignments on the opposite strand.

### Annotation of MusD full-length, genomic copies

All ERVK:ETnERV-int entries larger than 6 kb were collected from the current UCSC Golden Path repeat mask annotation (see above) as putative MusD full-length elements. Using *BEDTools closest*, upstream and downstream annotations on the same strand were collected to identify flanking LTR sequences and related sequences of the ETn superfamily that are contained between their flanking LTRs. Primer binding site (PBS) sequences were identified by a genome-wide alignment of tRF3a sequences (analogous to above described tRF3b alignments) and required to be immediately downstream of the 5’-LTR of elements larger than 6 kb. PBS coordinates were extended for 4 nt downstream using *BEDTools* (bedtools slop -s -l 4 -r 0) and converted to fasta format to collapse and count occurrences (Figure 5J). 71 elements matched all the above criteria. 65 are flanked by ERVK:ERVB7_1-LTR_MM, two by ERVK:ERVB7_2-LTR_MM, two by ERVK:ERVB7_2B-LTR_MM, and one each by ERVK:RLTR9E and ERVB7_4-LTR_MM. NCBI *ORFfinder* and *orfipy* (Singh & Wurtele, 2021) were used to find candidate open reading frames (ORFs) for all assembled MusD elements. 8 elements had all three retroviral ORFs (Gag 593 aa, Pr 318 aa, Pol 868 aa); the remaining elements had either truncated Pr or Pol ORFs. A total of 16 elements had intact Gag ORFs.

### MPRA experimental design and library construction

A massively parallel reporter assay (MPRA) was employed to test the effect of the tRNA primer binding site (PBS) sequence on MusD6 transposon expression. A pool of ∼1,700 single-stranded DNA oligos (*Twist Biosciences*) containing mutations at the PBS site and 4 nt downstream was cloned into pGL4.21-MusD6 (Schorn *et al*., 2017). In brief, the oligo pool was amplified using primer MPRA_#1 and MPRA_#2, *KAPA HiFi HotStart Polymerase* (Roche, # KK2502), and low cycle numbers to maintain sequence complexity and generate double-stranded DNA. The resulting PCR product was purified using *AMPure XP beads* (Beckman Coulter, #A63880) bead-to-sample ratio of 1.8X) and verified for the expected size using the *High Sensitivity DNA Kit* (Agilent #5067-4626). The pGL4.21-MusD6 plasmid backbone was linearized by restriction digest and amplified with *Q5 High Fidelity 2X Master Mix* (NEB, #M0491L) to generate a 20 bp overlap with the insert pool that was bulk cloned into the plasmid backbone using *NEBuilder HiFi DNA Assembly* (NEB #E5520). The assembled plasmid pool was electroporated into *10-beta electrocompetent E. coli* (NEB, #C3020K) in three independent transformations that resulted in replicate DNA input pools (MPRA DNA pool #1, #2, #3).

HeLa cells were seeded at 2x 10^5^ cells per 6-well one day prior to transfection with three different amounts (250 ng, 500 ng, 1500 ng) of DNA pool #1-3 using *Lipofectamine2000* (Thermo Fisher Scientific, #11668019), resulting in three (amounts) times three (DNA input pool), a total of nine conditions. One day post transfection, cells were washed with 1x phosphate buffered saline (Thermo Fisher Scientific, *#*20012050) and RNA was extracted using *TRIzol Reagent* (Thermo Fisher Scientific, #15596026). A total of 5 μg of DNase-treated RNA was reverse transcribed using *SuperScript IV VILO Master Mix* (Thermo Fisher Scientific, #11756050) which includes both, oligo(dT) and random primers. The resulting cDNA and the input plasmid pool DNA were amplified in nested PCR reactions first using primers MPRA_#3 and MPRA_#4, then using Illumina-P5-universal-for and Illumina-P7-index-rev primers. *Q5 High Fidelity 2X Master Mix* (NEB, #M0491L) was used for PCR amplification, and products were purified using a right-sided size selection with *AMPure XP beads* (Beckman Coulter, #A63880) using an initial bead-to-sample ratio of 0.7X followed by 1.1X, and eluted in *UltraPure Water* (Thermo Fisher Scientific, #10977049). Final libraries were verified for their expected size of ∼300 bp (*Agilent High Sensitivity DNA Kit,* Agilent #5067-4626), and quantified (*KAPA Library Quantification Kit Illumina platforms*, Roche #KR0405) before pooling. RNA-libraries (expression) and DNA-libraries (input plasmid pools) were sequenced on an *Illumina NextSeq500 Mid Output* single-end 150bp run. All oligo and primer sequences are listed in Supplemental Table S1.

### MPRA analysis

Reads were required to have 100% match with the upstream and downstream 5’-UTR sequence of MusD6 (Genbank AC124426, (Ribet *et al*, 2007)): (1) cutadapt -m 77 --error-rate 0 –overlap 48 -a ACGCTGCTTTGGAGCTCCACAGGAAAGGATCTTCGTATCGGACATCGGAGCA; (2) cutadapt --action none -m 48 --error-rate 0 --overlap 51 -g CCTTCTCAGTCGAACCGTTCACGTTGCGAGCTGCTGGCGGCCGCAACATTT. Subsequently, reads were required to align with zero mismatches to an index of the *Twist Biosciences* DNA sequence input pool using *Bowtie* (Langmead *et al*., 2009). *Bowtie2* (bowtie2 -N 1 -L 4 --rdg 1,1 --mp 2) (Langmead & Salzberg, 2012) and *Samtools* (samtools calmd -e) (Li *et al*, 2009) were used to generate CIGAR strings and to annotate mismatches and indels compared to the reference sequence. The reference sequence was the canonical PBS with a perfect 18 nt match with its Lysine^3^-TTT primer tRNA, extended for 4 more nucleotides of the MusD6 sequence, with the mismatch to m^1^A underlined: 5’-TGGCGCCCGAACAGGGAC<u>C</u>TGA-3’. Downstream analysis in this study only contains sequences without indels. Reads that fulfilled all above criteria were filtered for sequences that have mutations only within the 22 nt tRF3b target site (= 18 nt PBS plus 4 nt downstream), then clipped for all nucleotides upstream and downstream. Read counts were collected for sequences that had a minimum of 100 reads across all three DNA input libraries.

Relative abundances for each sequence in the MPRA library were calculated by dividing the read count of this sequence by the total read count in each condition. The original DNA library used for transfection was sequenced to establish baseline abundances. RNA/DNA fold changes were calculated by dividing RNA abundances by DNA abundances and log_10_-transformed. De-repression relative to the reference, canonical PBS was calculated by subtracting the log_10_-fold change of the reference sequence from each variant.

For the analysis of mutational effects by region or sliding window, we selected sequences containing mutations only in the region or sliding window of interest. For regional analysis, sequences were further stratified by mutation count within the region. Regions were defined as follows: “seed” region positions t2-7, “central” region positions t9-13, and “3’-region” positions t14-22. We used a one-sample t-test to assess the significance of mean de-repression relative to wild type. False discovery rates were corrected for multiple comparisons using the Benjamini-Hochberg method (Benjamini & Hochberg, 1995). All statistical analyses were performed using SciPy (Virtanen *et al*, 2020). For the analysis of expected effects of individual mutations, we used global epistasis regression with an additive genotype to latent phenotype model. This approach accounts for non-linear effects of multiple mutations due to global non-linearities in the measurement process. The analysis was performed using MAVE-NN (Tareen *et al*., 2022) with default parameters. To fit the global epistasis model, we randomly split the library into training (80%) and validation (20%) sets. The training set was used to fit the model while the validation set was used to evaluate model performance. Additive model parameters have a direct interpretation as the expected effects of mutations on the latent phenotype. However, interpretation of these effects is complicated by gauge freedoms in the model that are not constrained by the data. We resolved these gauge freedoms by constraining parameters corresponding to the canonical reference PBS sequence to zero, effectively using it as a reference point. Thus, parameter values reflect effects on the latent phenotype relative to the canonical PBS sequence.

### Luciferase assays

A number of sequences from the MPRA plasmid pool were cloned as single sequences and tested for luciferase expression. Oligos (Supplemental Table S1) were amplified with primers L1/L2 (Supplemental Table S1) by 14 cycles of PCR with *KOD Xtreme Hot Start DNA Polymerase* (Sigma-Aldrich, #71975-M). The resulting dsDNA was purified using *RNAClean XP beads* (Beckman Coulter, #A63987). The pGL4.21-MusD6 plasmid (Schorn *et al*., 2017) was amplified by 30 cycles of PCR with *KOD Xtreme Hot Start DNA Polymerase* (Sigma-Aldrich, #71975-M) and primers L3/L4 (Supplemental Table S1), and purified by agarose gel extraction. The pool of dsDNA was inserted into the linearized vector by *NEBuilder HiFi DNA Assembly* (NEB #E5520). Individual colonies were picked to isolate DNA and sequence confirmed by Sanger sequencing (primer L5, Supplemental Table S1) and *Plasmidsaurus* whole plasmid sequencing. 5x 10^4^ HeLa cells were transfected in 24-well plates with 5 ng of each pGL4.21 *Firefly* luciferase reporter constructs, and 205 ng of pcDNA3.1(+)-neo as filler DNA per well using *Lipofectamine2000* (Thermo Fisher Scientific, #11668019). Cells were lysed 24 h post transfection in 100 ul of Glo Lysis buffer (Promega, #E2661). 40 ul of lysate per sample were used to measure luciferase activity using the *Dual-Glo Luciferase Assay System* (Promega, #E2940) according to the manufacturer’s instructions, using a *GloMax Discover Microplate reader* (Promega) at 0.3 sec integration time. Relative light units (RLU) were calculated by normalizing *Firefly* luminescence values to total protein amount determined by Bradford Protein Assay (BioRad, #5000006), measuring absorbance at 595 nm.

### Statistical analysis

*PRISM* version 10.3.1 was used to perform statistical analyses of luciferase and transposition assay results using two-tailed Student’s t-test. *R* version 4.4.1 was used for data visualization and statistical analysis of read counts in *DESeq2* (Love *et al*., 2014). Software for sequence analysis was installed and updated using *Bioconda* (Grüning *et al*, 2018). For statistical analysis of the massively parallel reporter assay data, see section ’MPRA analysis’.

---

**Extended Data Figure 1.**
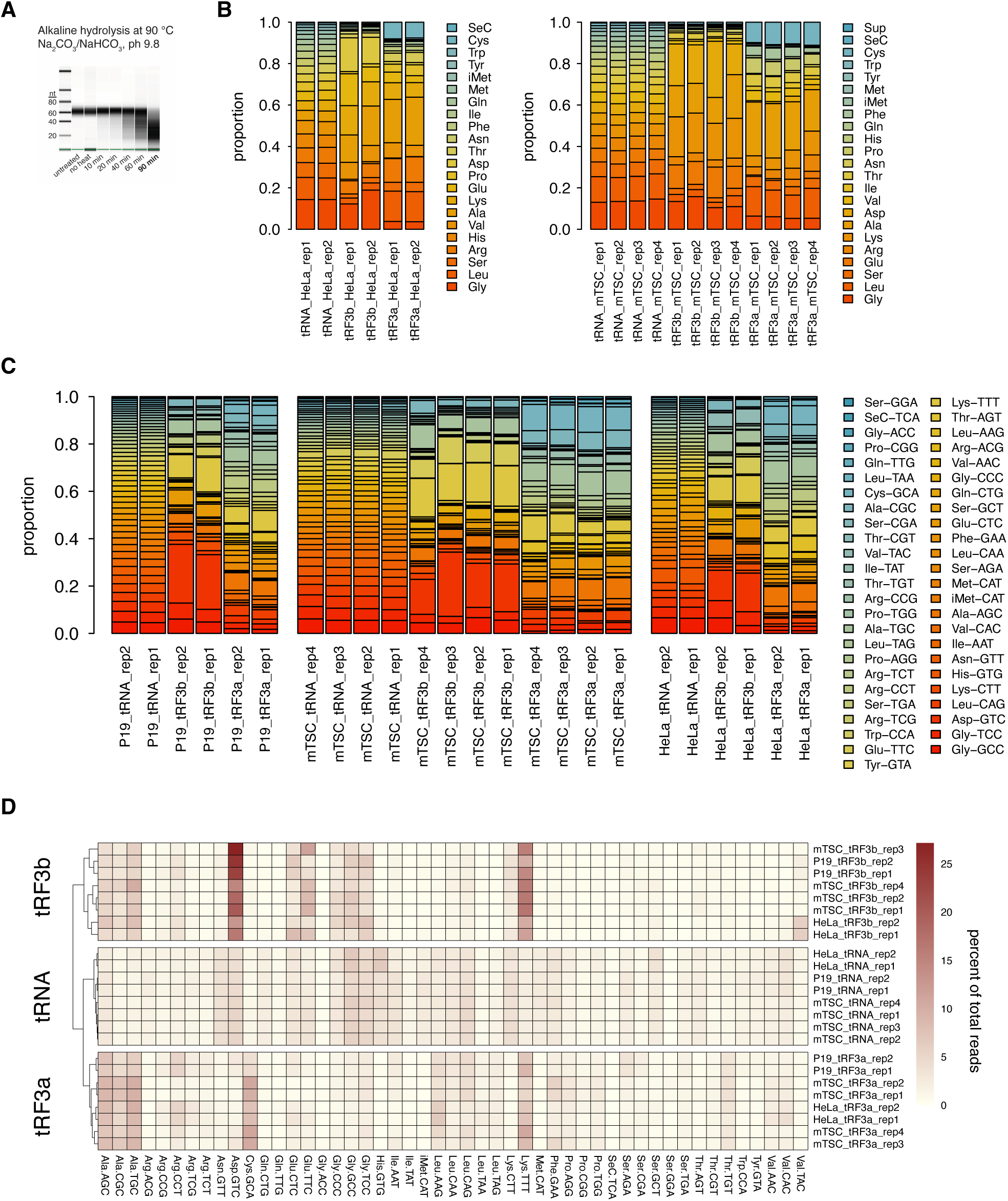
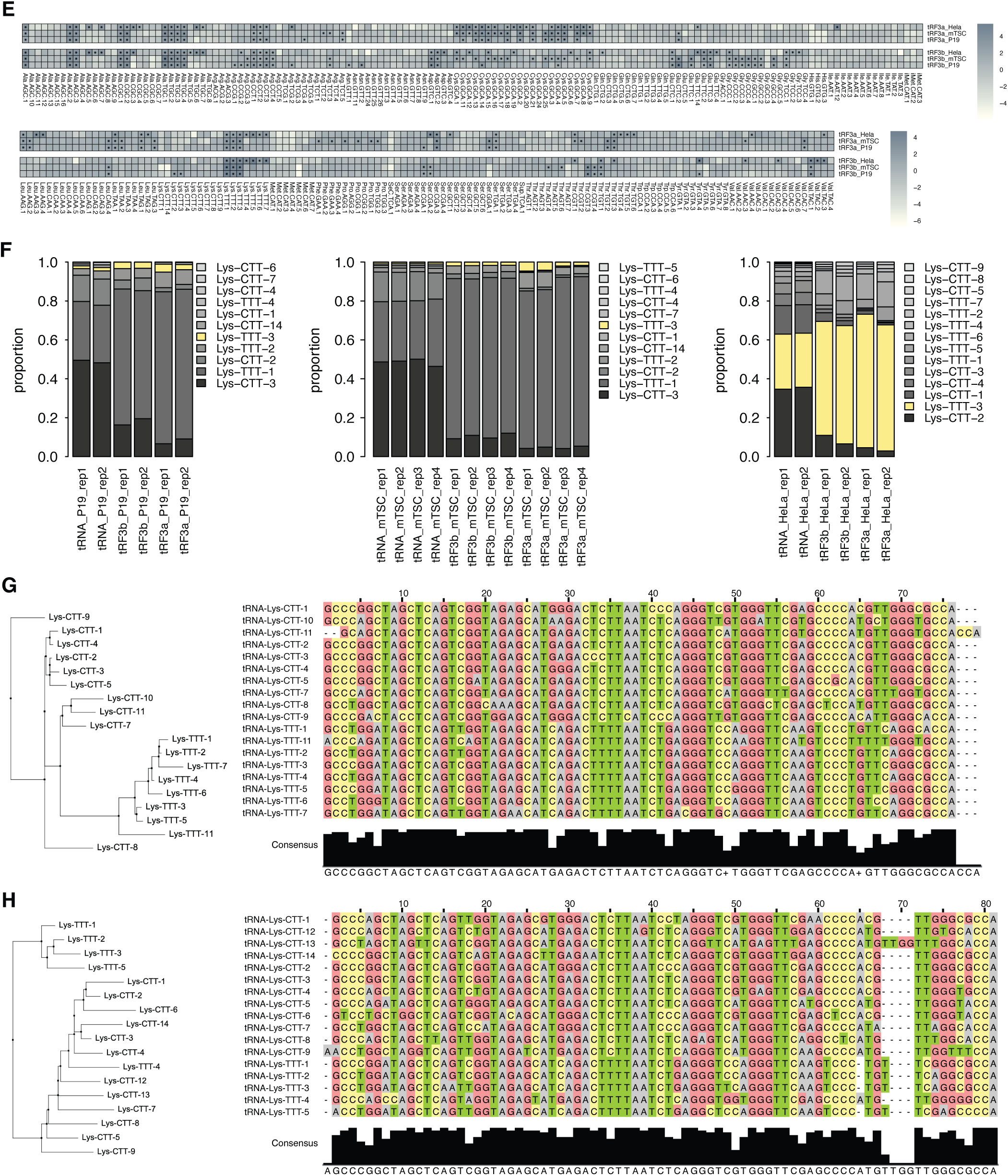
3’-tRFs and full-length tRNAs in mouse and human cell lines. **(A)** Optimization of alkaline hydrolysis conditions for tRNA hydrolysis sequencing (Gogakos *et al*., 2017). **(B)** Relates to Figure 1C: Proportion of full-length tRNAs, tRF3b and tRF3a fragments by amino acid isotype in HeLa cells (two biological replicates) and mTSC (four biological replicates). **(C)** Proportion of tRNA, tRF3a, and tRF3b isoacceptor sequences in all three cell lines (P19, mTSC, HeLa) and their biological replicates. **(D)** tRNA, tRF3a, and tRF3b expression in percent of total reads clustered by similarity for all cell lines. **(E)** Relates to Figure 1F: Heat map of log_2_-fold changes (log2FC) of tRF3a and tRF3b relative to their corresponding full-length tRNAs for all isodecoder sequences. Asterisks (*) mark significant enrichment (log2FC >1, adjusted p-value < 0.01). **(F)** Proportion of Lysine isodecoder tRNA, tRF3a, and tRF3b sequences for all three cell lines (P19, mTSC, HeLa). Lys-TTT-3 (yellow) also referred to as Lys^3^-TTT is the primer tRNA for MusD/ETn. **(G)** Phylogenetic tree of human Lysine tRNA isodecoder sequences and Clustal Omega alignment of the 3’-end of cytoplasmic isodecoder sequences. **(H)** Phylogenetic tree of mouse Lysine tRNA isodecoder sequences and Clustal Omega alignment of the 3’-end of cytoplasmic isodecoder sequences.

**Extended Data Figure 2.**
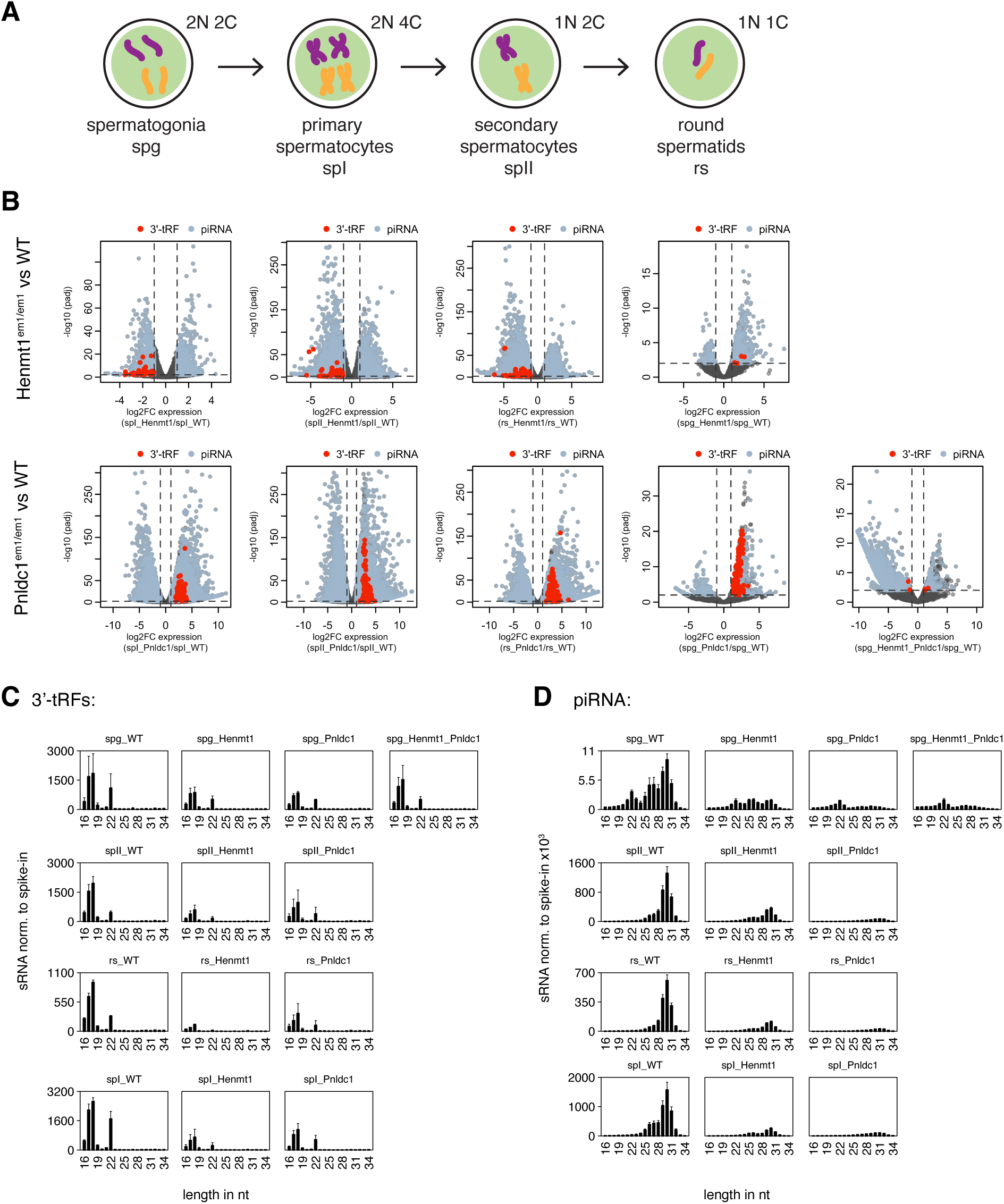
3’-tRF analysis of small RNAs from male germ cells of *Henmt1^em1/em1^, Pnldc1^em1/em1^,* and wildtype mice (Gainetdinov *et al*., 2021) **(A)** Cartoon of male germ cell types that were FACS-sorted for RNA isolation and small RNA sequencing by (Gainetdinov *et al*., 2021). Illustration modified from (Gainetdinov *et al*, 2018). **(B)** Relates to Figure 2E: Volcano plot showing log_2_-fold changes (log2FC) of small RNAs in *Henmt1^em1/em1^* and *Pnldc1^em1/em1^* germ cells compared to wildtype. Each dot represents one sequence (piRNA, miRNA) or isoacceptor (3’-tRFs). Significantly enriched or depleted (Log2FC > |1| and padj. < 0.01) piRNAs and 3’-tRFs are colored in steel-blue and red, respectively. **(C)** Relates to Figure 2D: Size distribution of 3’-tRFs in male germ cells (abbreviations see panel A) from wildtype *versus* loss of function *Henmt1* and *Plndc1* mice. **(D)** Relates to Figure 2D: Size distribution of piRNAs in male germ cells (abbreviations see panel A) from wildtype *versus* loss of function *Henmt1* and *Plndc1* mice.

**Extended Data Figure 3.**
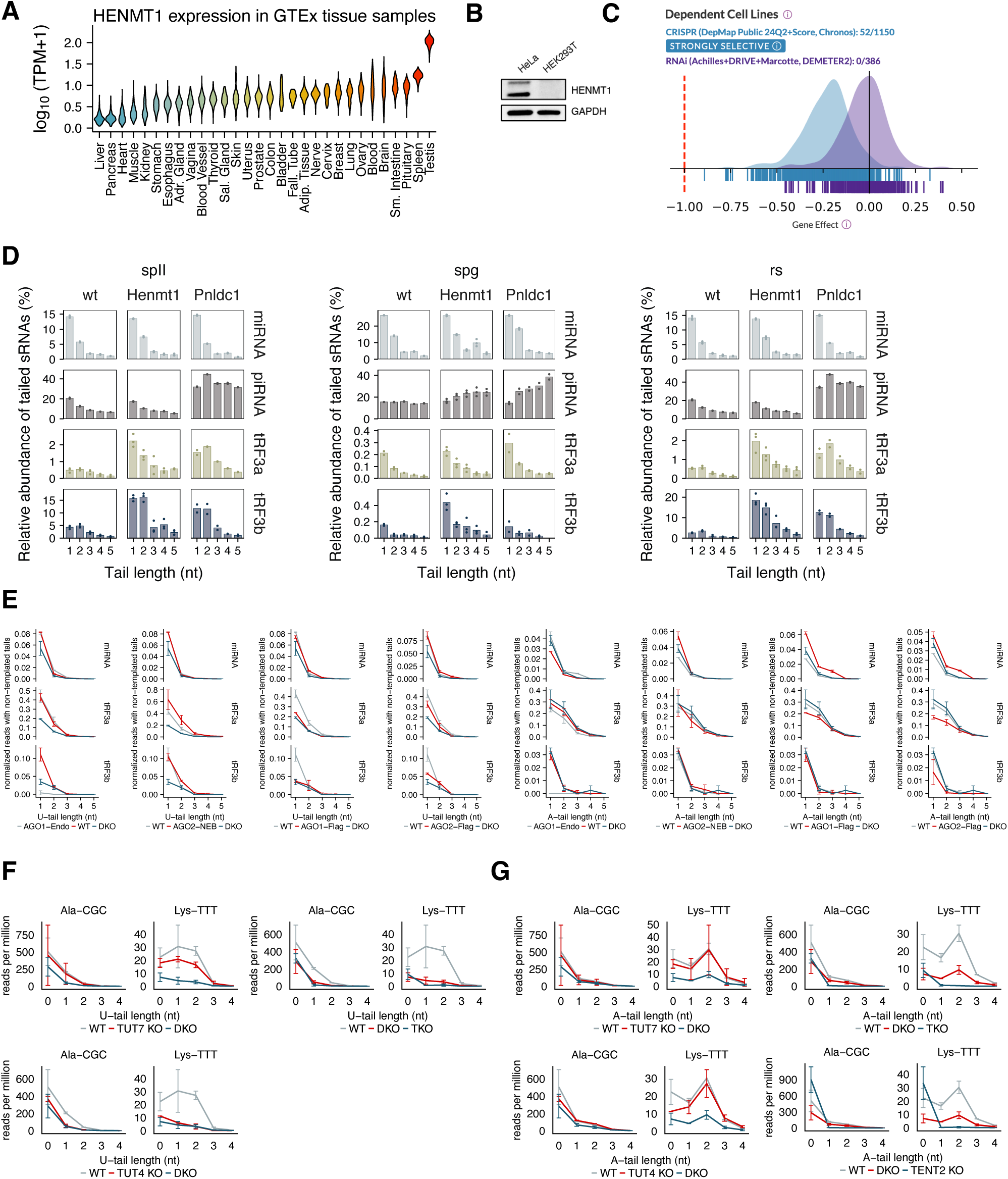
HENMT1 expression outside the germline and 3’-tRF untemplated nucleotide tailing. **(A)** *HENMT1* RNA expression in human GTEx tissue samples. **(B)** Western blot of lysates from HEK293T and HeLa cells probing for HENMT1 expression. **(C)** DepMap ’Gene Effect’ score of HENMT1 knockdown (RNAi) and knockout (CRISPR) **(D)** Length of untemplated 3’-nucleotide tails in germ cell types of *Henmt1^em1/em1^, Pnldc1^em1/em1^,* and wildtype mice [data Gainetdinov et al. 2021]. **(E)** Uridine and adenosine tail length in AGO immunoprecipitated (IP) and total RNA (designated “WT”) of HEK293T cells. DKO: double knockout of TUT4 and TUT7; AGO1-Endo: pulldown of endogenous AGO1; AGO2-NEB: pulldown of AGO2 with New England Biolabs (NEB) antibody; AGO1/AGO2-Flag: FLAG-tagged, overexpressed AGO1/AGO2 **(F)** Uridine and **(G)** adenosine tail length of AGO2-bound Ala-CGC and Lys-TTT isoacceptor sequences in HEK293T wildtype (WT) and nucleotidyltransferase KO cells. DKO: double-knockout of TUT4 and TUT7; TKO: TUT4/TUT7/TENT2 triple-knockout.

**Extended Data Figure 4.**
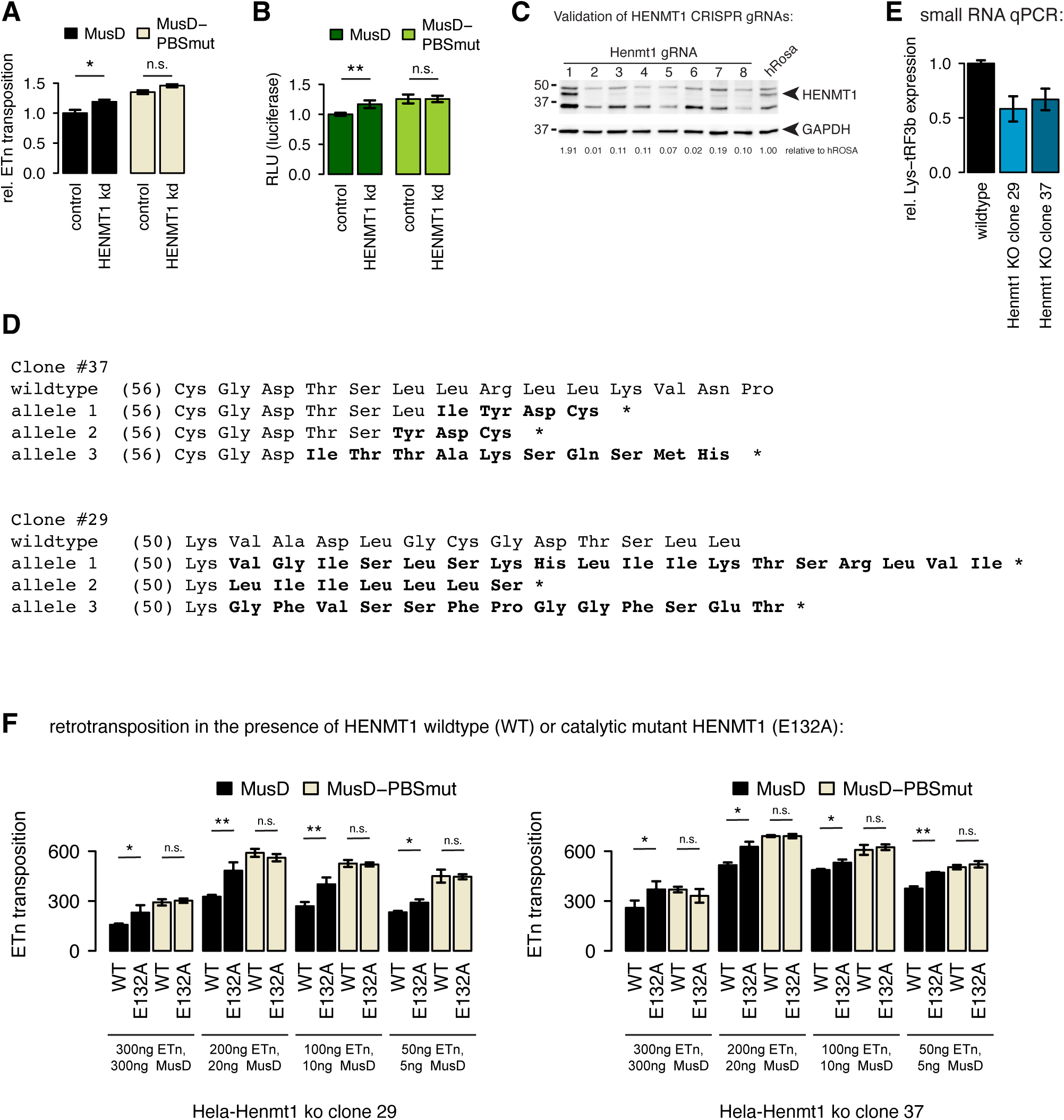
Retrotransposition and small RNA expression during HENMT1 knockdown and in *HENMT1* KO clonal cell lines. **(A)** Plasmid-based retrotransposition assay during HENMT1 knockdown using *onTarget-plus siRNAs* (Dharmacon). **(B)** Luciferase expression from reporter constructs containing the MusD 5’-UTR with wildtype and mutated PBS (Schorn *et al*., 2017) during knockdown of HENMT1. The luciferase ORF is exactly in place of the first MusD ORF. RLU, relative light units. **(C)** Western Blot validation of HENMT1 expression in CRISPR-Cas9 HeLa cells using different guide RNAs (gRNA). Guides targeting the human Rosa (hRosa) locus were used as a negative control. All guides except gRNA #1 showed decrease of HENMT1 expression. Protein amounts were estimated probing the same membrane for GAPDH expression. **(D)** Sanger-sequencing traces of genomic DNA isolated from HeLa-Cas9 *HENMT1*-g2 clonal cell lines 29 and 37 was resolved by CRISPR-ID (Dehairs *et al*, 2016) to show mutations in all alleles of polyploid cells. The resulting amino acid sequences and premature termination (*) are indicated. Guide 2 targets between Cys-56 and Arg-63, at the beginning of the methyltransferase domain. Numbers in brackets indicate amino acid position. **(E)** Small RNA quantitative PCR (*Taqman miRNA Advanced Assay*, Thermo Fisher Scientific) with custom primer-probes to detect Lys^3^-tRF3b in HeLa-Cas9 wildtype progenitor cells and clonal *HENMT1* knockout (KO) cells. Relative expression was normalized to miR-191-5p. Values are mean expression +/- standard error. **(F)** Retrotransposition assays in clonal *HENMT1* CRISPR KO HeLa cell lines, clone #29 and #37, with transient expression of functional, wildtype *HENMT1* or the catalytic mutant *HENMT1-E132A* in a pcDNA3.1 vector backbone. Depicted are colony counts when using different amounts of MusD wildtype or PBS^mut^ plus ETn-reporter plasmid. Loss of HENMT1 methyltransferase activity increased MusD-mediated ETn retrotransposition dependent on the wildtype MusD PBS sequence. Values are the mean of colony counts +/- standard deviation.

**Extended Data Figure 5.**
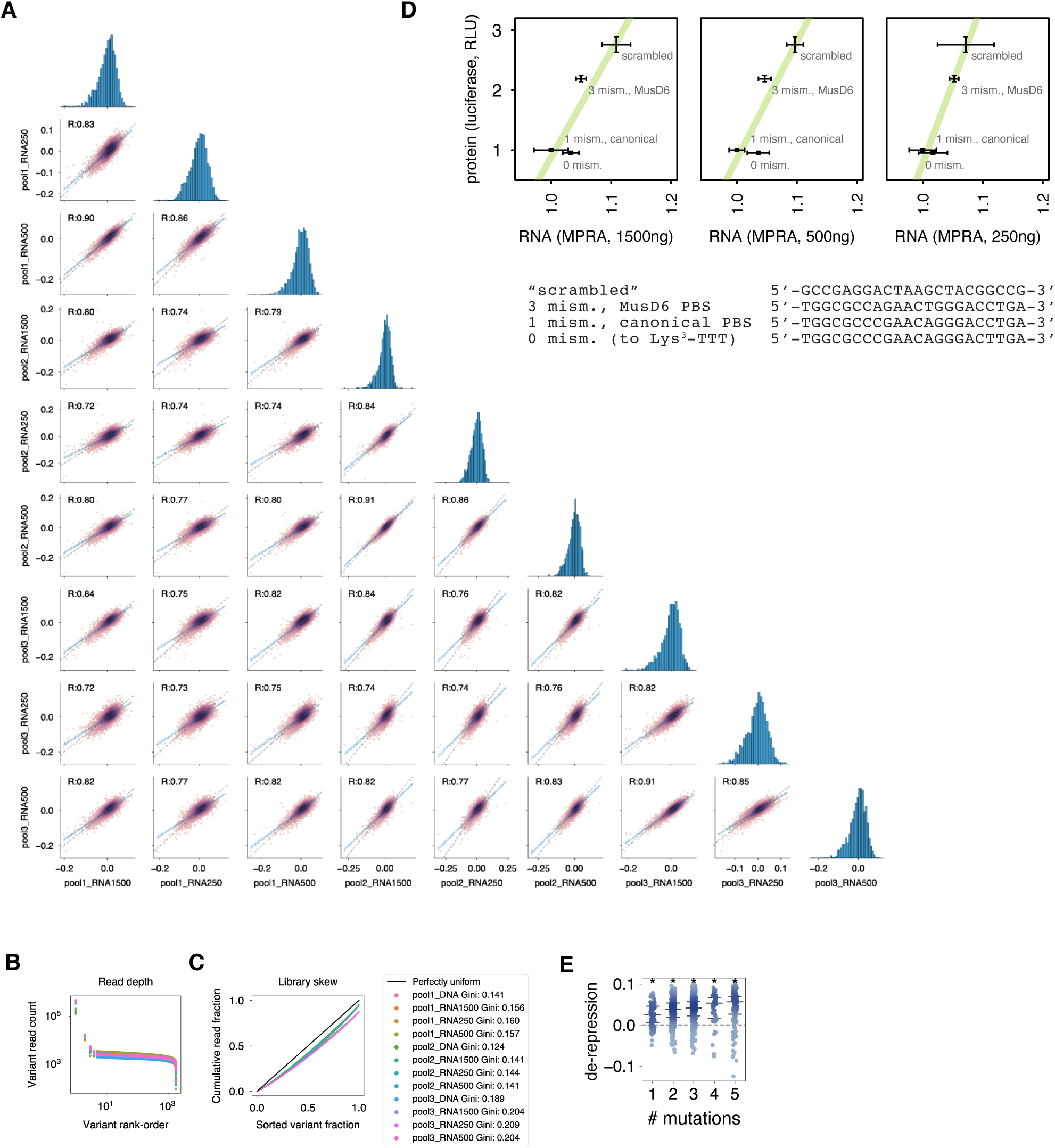
Validation of MPRA. **(A)** Replicate correspondence plots. Each point represents the log_10_-fold change between RNA and DNA abundance of an individual MPRA sequence as measured in two different MPRA replicates. Grey dotted lines indicate equality. Blue dashed lines represented a linear fit of the relationship between the observed values in the two replicates. Pearson R values representing correlation between that pair of replicates is included in each panel. Histograms represent the distribution of fitness values observed in each replicate. **(B)** Zipf plot for diagnosing over or under abundant sequences in the MPRA libraries. Each point is the number of reads observed for one MPRA sequence sorted in descending order of read count. Most sequences have similar numbers of read counts but a small number have very high or very low count numbers. **(C)** Lorenz plots for diagnosing inequality in library read counts. Each point the fraction of total library reads accounted for by MPRA sequences in the bottom x fraction of the library (when sorted in ascending order by read count). The black line indicates what a library with perfectly uniform read counts would look like. Deviations from perfect equality are quantified using the Gini index which represents the fractional difference in the area under each library Lorenz curve and the uniform Lorenz curve. Values close to 0 are highly uniform and values close to 1 are highly skewed. **(D)** Plots show RNA expression of select luciferase reporter plasmids quantified in the MPRA and their protein expression quantified by luciferase assays and expressed in relative light units (RLU). Expression was normalized to the values of the canonical reference PBS sequence described in the main text (Figure 5). **(E)** Observed de-repression dependent on the number of mutations relative to the canonical, reference PBS sequence. Stars indicate significant de-repression (Benjamini-Hochberg-adjusted p<0.05, 1-sample t-test). Lines represent median and quartiles and all data points are plotted.

**Supplementary Table S1.**
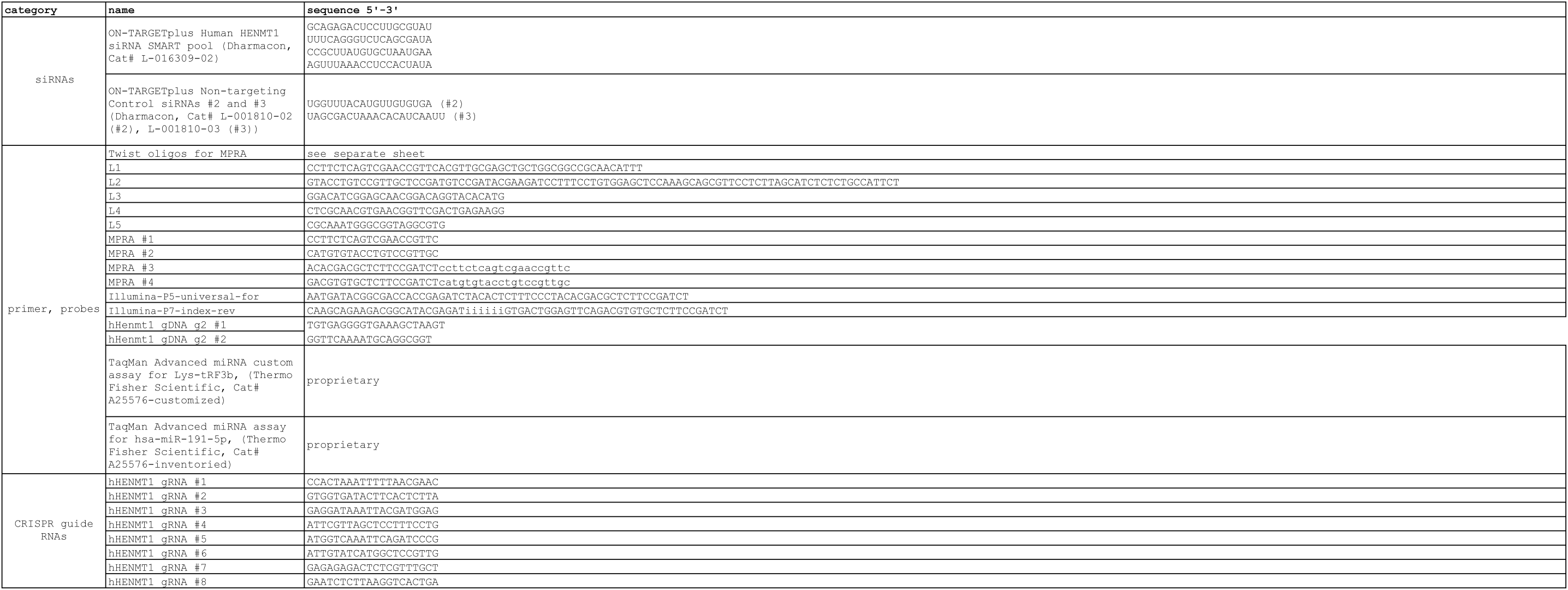
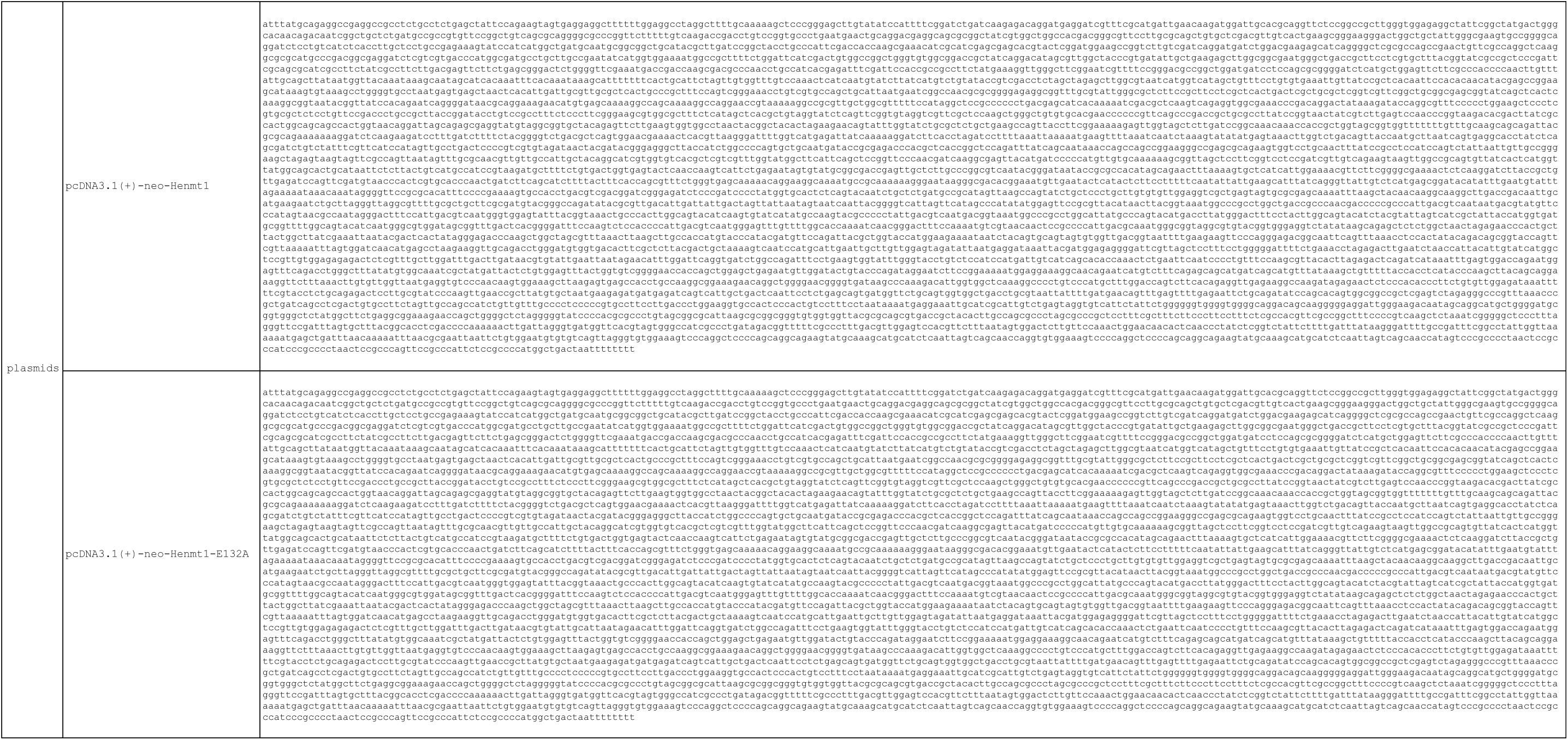

## Notes

### Competing Interest Statement

The authors have declared no competing interest.

### Summary of Updates

Fixed bibliography order and extended analysis for figure 1.

## REFERENCES

1. Anzelon TA, Chowdhury S, Hughes SM, Xiao Y, Lander GC, MacRae IJ (2021) Structural basis for piRNA targeting. Nature 597: 285–289

2. Barnett DW, Garrison EK, Quinlan AR, Stromberg MP, Marth GT (2011) BamTools: a C++ API and toolkit for analyzing and managing BAM files. Bioinformatics 27: 1691–1692

3. Bartel DP (2009) MicroRNAs: target recognition and regulatory functions. Cell 136: 215–233

4. Begik O, Lucas MC, Liu H, Ramirez JM, Mattick JS, Novoa EM (2020) Integrative analyses of the RNA modification machinery reveal tissue- and cancer-specific signatures. Genome Biol 21: 97

5. Benjamini Y, Hochberg Y (1995) Controlling the false discovery rate: a practical and powerful approach to multiple testing. Journal of the Royal Statistical Society: Series B (Methodological) 57: 289–300

6. Berwin B, Barklis E (1993) Retrovirus-mediated insertion of expressed and non-expressed genes at identical chromosomal locations. Nucleic Acids Res 21: 2399–2407

7. Carithers LJ, Ardlie K, Barcus M, Branton PA, Britton A, Buia SA, Compton CC, DeLuca DS, Peter-Demchok J, Gelfand ET et al (2015) A Novel Approach to High-Quality Postmortem Tissue Procurement: The GTEx Project. Biopreserv Biobank 13: 311–319

8. Chapman KB, Byström AS, Boeke JD (1992) Initiator methionine tRNA is essential for Ty1 transposition in yeast. Proc Natl Acad Sci U S A 89: 3236–3240

9. Couvillion MT, Sachidanandam R, Collins K (2010) A growth-essential Tetrahymena Piwi protein carries tRNA fragment cargo. Genes Dev 24: 2742–2747

10. Cullen H, Schorn AJ (2020) Endogenous Retroviruses Walk a Fine Line between Priming and Silencing. Viruses 12

11. Das AT, Klaver B, Berkhout B (1997) Sequence variation of the human immunodeficiency virus primer-binding site suggests the use of an alternative tRNA(Lys) molecule in reverse transcription. J Gen Virol 78 (Pt 4): 837–840

12. Dehairs J, Talebi A, Cherifi Y, Swinnen JV (2016) CRISP-ID: decoding CRISPR mediated indels by Sanger sequencing. Sci Rep 6: 28973

13. Enright AJ, John B, Gaul U, Tuschl T, Sander C, Marks DS (2003) MicroRNA targets in Drosophila. Genome Biol 5: R1

14. Faulkner GJ, Kimura Y, Daub CO, Wani S, Plessy C, Irvine KM, Schroder K, Cloonan N, Steptoe AL, Lassmann T et al (2009) The regulated retrotransposon transcriptome of mammalian cells. Nat Genet 41: 563–571

15. Fennessey CM, Camus C, Immonen TT, Reid C, Maldarelli F, Lifson JD, Keele BF (2019) Low-level alternative tRNA priming of reverse transcription of HIV-1 and SIV in vivo. Retrovirology 16: 11

16. Flemr M, Malik R, Franke V, Nejepinska J, Sedlacek R, Vlahovicek K, Svoboda P (2013) A retrotransposon-driven dicer isoform directs endogenous small interfering RNA production in mouse oocytes. Cell 155: 807–816

17. Franke V, Ganesh S, Karlic R, Malik R, Pasulka J, Horvat F, Kuzman M, Fulka H, Cernohorska M, Urbanova J et al (2017) Long terminal repeats power evolution of genes and gene expression programs in mammalian oocytes and zygotes. Genome Res 27: 1384–1394

18. Gagnier L, Belancio VP, Mager DL (2019) Mouse germ line mutations due to retrotransposon insertions. Mob DNA 10: 15

19. Gainetdinov I, Colpan C, Arif A, Cecchini K, Zamore PD (2018) A Single Mechanism of Biogenesis, Initiated and Directed by PIWI Proteins, Explains piRNA Production in Most Animals. Mol Cell 71: 775–790.e775

20. Gainetdinov I, Colpan C, Cecchini K, Albosta P, Jouravleva K, Vega-Badillo J, Lee Y, Özata DM, Zamore PD (2020) Terminal Modification, Sequence, and Length Determine Small RNA Stability in Animals. bioRxiv: 2020.2009.2008.287979

21. Gainetdinov I, Colpan C, Cecchini K, Arif A, Jouravleva K, Albosta P, Vega-Badillo J, Lee Y, Özata DM, Zamore PD (2021) Terminal modification, sequence, length, and PIWI-protein identity determine piRNA stability. Mol Cell 81: 4826–4842.e4828

22. Gainetdinov I, Vega-Badillo J, Cecchini K, Bagci A, Colpan C, De D, Bailey S, Arif A, Wu PH, MacRae IJ et al (2023) Relaxed targeting rules help PIWI proteins silence transposons. Nature 619: 394–402

23. Gao L, Behrens A, Rodschinka G, Forcelloni S, Wani S, Strasser K, Nedialkova DD (2024) Selective gene expression maintains human tRNA anticodon pools during differentiation. Nat Cell Biol 26: 100–112

24. Giraldez MD, Spengler RM, Etheridge A, Godoy PM, Barczak AJ, Srinivasan S, De Hoff PL, Tanriverdi K, Courtright A, Lu S et al (2018) Comprehensive multi-center assessment of small RNA-seq methods for quantitative miRNA profiling. Nat Biotechnol 36: 746–757

25. Gogakos T, Brown M, Garzia A, Meyer C, Hafner M, Tuschl T (2017) Characterizing Expression and Processing of Precursor and Mature Human tRNAs by Hydro-tRNAseq and PAR-CLIP. Cell Rep 20: 1463–1475

26. Goodarzi H, Nguyen HCB, Zhang S, Dill BD, Molina H, Tavazoie SF (2016) Modulated Expression of Specific tRNAs Drives Gene Expression and Cancer Progression. Cell 165: 1416–1427

27. Grüning B, Dale R, Sjödin A, Chapman BA, Rowe J, Tomkins-Tinch CH, Valieris R, Köster J (2018) Bioconda: sustainable and comprehensive software distribution for the life sciences. Nat Methods 15: 475–476

28. Guo Y, Li TD, Modzelewski AJ, Siomi H (2024) Retrotransposon renaissance in early embryos. Trends Genet 40: 39–51

29. Han J, Mendell JT (2023) MicroRNA turnover: a tale of tailing, trimming, and targets. Trends Biochem Sci 48: 26–39

30. Hasler D, Lehmann G, Murakawa Y, Klironomos F, Jakob L, Grasser FA, Rajewsky N, Landthaler M, Meister G (2016) The Lupus Autoantigen La Prevents Mis-channeling of tRNA Fragments into the Human MicroRNA Pathway. Mol Cell 63: 110–124

31. Haussecker D, Huang Y, Lau A, Parameswaran P, Fire AZ, Kay MA (2010) Human tRNA-derived small RNAs in the global regulation of RNA silencing. RNA 16: 673–695

32. Herridge RP, Dolata J, Migliori V, de Santis Alves C, Borges F, Schorn AJ, van Ex F, Lin A, Bajczyk M, Parent JS et al (2025) Pseudouridine guides germline small RNA transport and epigenetic inheritance. Nat Struct Mol Biol 32: 277–286

33. Horwich MD, Li C, Matranga C, Vagin V, Farley G, Wang P, Zamore PD (2007) The Drosophila RNA methyltransferase, DmHen1, modifies germline piRNAs and single-stranded siRNAs in RISC. Curr Biol 17: 1265–1272

34. Hu WS, Hughes SH (2012) HIV-1 reverse transcription. Cold Spring Harb Perspect Med 2

35. Jansz N, Faulkner GJ (2021) Endogenous retroviruses in the origins and treatment of cancer. Genome Biol 22: 147

36. Ji L, Chen X (2012) Regulation of small RNA stability: methylation and beyond. Cell Res 22: 624–636

37. Kamminga LM, Luteijn MJ, den Broeder MJ, Redl S, Kaaij LJ, Roovers EF, Ladurner P, Berezikov E, Ketting RF (2010) Hen1 is required for oocyte development and piRNA stability in zebrafish. Embo j 29: 3688–3700

38. Kawaji H, Nakamura M, Takahashi Y, Sandelin A, Katayama S, Fukuda S, Daub CO, Kai C, Kawai J, Yasuda J et al (2008) Hidden layers of human small RNAs. BMC Genomics 9: 157

39. Keeney JB, Chapman KB, Lauermann V, Voytas DF, Astrom SU, von Pawel-Rammingen U, Bystrom A, Boeke JD (1995) Multiple molecular determinants for retrotransposition in a primer tRNA. Mol Cell Biol 15: 217–226

40. Kinney JB, McCandlish DM (2019) Massively Parallel Assays and Quantitative Sequence-Function Relationships. Annu Rev Genomics Hum Genet 20: 99–127

41. Kirino Y, Mourelatos Z (2007) Mouse Piwi-interacting RNAs are 2’-O-methylated at their 3’ termini. Nat Struct Mol Biol 14: 347–348

42. Kozomara A, Birgaoanu M, Griffiths-Jones S (2019) miRBase: from microRNA sequences to function. Nucleic Acids Res 47: D155–d162

43. Kumar P, Anaya J, Mudunuri SB, Dutta A (2014) Meta-analysis of tRNA derived RNA fragments reveals that they are evolutionarily conserved and associate with AGO proteins to recognize specific RNA targets. BMC Biol 12: 78

44. Kurth HM, Mochizuki K (2009) 2’-O-methylation stabilizes Piwi-associated small RNAs and ensures DNA elimination in Tetrahymena. RNA 15: 675–685

45. Kuscu C, Kumar P, Kiran M, Su Z, Malik A, Dutta A (2018) tRNA fragments (tRFs) guide Ago to regulate gene expression post-transcriptionally in a Dicer-independent manner. Rna 24: 1093–1105

46. Langmead B, Salzberg SL (2012) Fast gapped-read alignment with Bowtie 2. Nat Methods 9: 357–359

47. Langmead B, Trapnell C, Pop M, Salzberg SL (2009) Ultrafast and memory-efficient alignment of short DNA sequences to the human genome. Genome Biol 10: R25

48. Le Grice SF (2003) ”In the beginning”: initiation of minus strand DNA synthesis in retroviruses and LTR-containing retrotransposons. Biochemistry 42: 14349–14355

49. Li H, Handsaker B, Wysoker A, Fennell T, Ruan J, Homer N, Marth G, Abecasis G, Durbin R, Genome Project Data Processing S (2009) The Sequence Alignment/Map format and SAMtools. Bioinformatics 25: 2078–2079

50. Li Z, Ender C, Meister G, Moore PS, Chang Y, John B (2012) Extensive terminal and asymmetric processing of small RNAs from rRNAs, snoRNAs, snRNAs, and tRNAs. Nucleic Acids Res 40: 6787–6799

51. Lim SL, Qu ZP, Kortschak RD, Lawrence DM, Geoghegan J, Hempfling AL, Bergmann M, Goodnow CC, Ormandy CJ, Wong L et al (2015) HENMT1 and piRNA Stability Are Required for Adult Male Germ Cell Transposon Repression and to Define the Spermatogenic Program in the Mouse. PLoS Genet 11: e1005620

52. Liu S, Harada BT, Miller JT, Le Grice SF, Zhuang X (2010) Initiation complex dynamics direct the transitions between distinct phases of early HIV reverse transcription. Nat Struct Mol Biol 17: 1453–1460

53. Liu X, Liu Z, Wu Z, Ren J, Fan Y, Sun L, Cao G, Niu Y, Zhang B, Ji Q et al (2023) Resurrection of endogenous retroviruses during aging reinforces senescence. Cell 186: 287–304.e226

54. Loubalova Z, Fulka H, Horvat F, Pasulka J, Malik R, Hirose M, Ogura A, Svoboda P (2021) Formation of spermatogonia and fertile oocytes in golden hamsters requires piRNAs. Nat Cell Biol 23: 992–1001

55. Love MI, Huber W, Anders S (2014) Moderated estimation of fold change and dispersion for RNA-seq data with DESeq2. Genome Biol 15: 550

56. Lv X, Xiao W, Lai Y, Zhang Z, Zhang H, Qiu C, Hou L, Chen Q, Wang D, Gao Y et al (2023) The non-redundant functions of PIWI family proteins in gametogenesis in golden hamsters. Nat Commun 14: 5267

57. Maksakova IA, Mager DL (2005) Transcriptional regulation of early transposon elements, an active family of mouse long terminal repeat retrotransposons. J Virol 79: 13865–13874

58. Markopoulos G, Noutsopoulos D, Mantziou S, Gerogiannis D, Thrasyvoulou S, Vartholomatos G, Kolettas E, Tzavaras T (2016) Genomic analysis of mouse VL30 retrotransposons. Mob DNA 7: 10

59. Martin M (2011) Cutadapt removes adapter sequences from high-throughput sequencing reads. 2011 17

60. Matsui T, Leung D, Miyashita H, Maksakova IA, Miyachi H, Kimura H, Tachibana M, Lorincz MC, Shinkai Y (2010) Proviral silencing in embryonic stem cells requires the histone methyltransferase ESET. Nature 464: 927–931

61. Maute RL, Schneider C, Sumazin P, Holmes A, Califano A, Basso K, Dalla-Favera R (2013) tRNA-derived microRNA modulates proliferation and the DNA damage response and is down-regulated in B cell lymphoma. Proc Natl Acad Sci U S A 110: 1404–1409

62. McGeary SE, Bisaria N, Pham TM, Wang PY, Bartel DP (2022) MicroRNA 3’-compensatory pairing occurs through two binding modes, with affinity shaped by nucleotide identity and position. Elife 11

63. McGeary SE, Lin KS, Shi CY, Pham TM, Bisaria N, Kelley GM, Bartel DP (2019) The biochemical basis of microRNA targeting efficacy. Science 366

64. Modepalli V, Fridrich A, Agron M, Moran Y (2018) The methyltransferase HEN1 is required in Nematostella vectensis for microRNA and piRNA stability as well as larval metamorphosis. PLoS Genet 14: e1007590

65. Modzelewski AJ, Shao W, Chen J, Lee A, Qi X, Noon M, Tjokro K, Sales G, Biton A, Anand A et al (2021) A mouse-specific retrotransposon drives a conserved Cdk2ap1 isoform essential for development. Cell 184: 5541–5558.e5522

66. Ohara T, Sakaguchi Y, Suzuki T, Ueda H, Miyauchi K, Suzuki T (2007) The 3’ termini of mouse Piwi-interacting RNAs are 2’-O-methylated. Nat Struct Mol Biol 14: 349–350

67. Parikh R, Wilson B, Marrah L, Su Z, Saha S, Kumar P, Huang F, Dutta A (2022) tRForest: a novel random forest-based algorithm for tRNA-derived fragment target prediction. NAR Genom Bioinform 4: lqac037

68. Park W, Li J, Song R, Messing J, Chen X (2002) CARPEL FACTORY, a Dicer homolog, and HEN1, a novel protein, act in microRNA metabolism in Arabidopsis thaliana. Curr Biol 12: 1484–1495

69. Pastore B, Hertz HL, Price IF, Tang W (2021) pre-piRNA trimming and 2’-O-methylation protect piRNAs from 3’ tailing and degradation in C. elegans. Cell Rep 36: 109640

70. Peaston AE, Evsikov AV, Graber JH, de Vries WN, Holbrook AE, Solter D, Knowles BB (2004) Retrotransposons regulate host genes in mouse oocytes and preimplantation embryos. Dev Cell 7: 597–606

71. Peng L, Zhang F, Shang R, Wang X, Chen J, Chou JJ, Ma J, Wu L, Huang Y (2018) Identification of substrates of the small RNA methyltransferase Hen1 in mouse spermatogonial stem cells and analysis of its methyl-transfer domain. J Biol Chem 293: 9981–9994

72. Quinlan AR, Hall IM (2010) BEDTools: a flexible suite of utilities for comparing genomic features. Bioinformatics 26: 841–842

73. Rebollo R, Romanish MT, Mager DL (2012) Transposable elements: an abundant and natural source of regulatory sequences for host genes. Annu Rev Genet 46: 21–42

74. Ribet D, Dewannieux M, Heidmann T (2004) An active murine transposon family pair: retrotransposition of “master” MusD copies and ETn trans-mobilization. Genome Res 14: 2261–2267

75. Ribet D, Harper F, Dewannieux M, Pierron G, Heidmann T (2007) Murine MusD retrotransposon: structure and molecular evolution of an “intracellularized” retrovirus. J Virol 81: 1888–1898

76. Rosenkranz D, Zischler H, Gebert D (2022) piRNAclusterDB 2.0: update and expansion of the piRNA cluster database. Nucleic Acids Res 50: D259–d264

77. Rowe HM, Jakobsson J, Mesnard D, Rougemont J, Reynard S, Aktas T, Maillard PV, Layard-Liesching H, Verp S, Marquis J et al (2010) KAP1 controls endogenous retroviruses in embryonic stem cells. Nature 463: 237–240

78. Ryvkin P, Leung YY, Silverman IM, Childress M, Valladares O, Dragomir I, Gregory BD, Wang LS (2013) HAMR: high-throughput annotation of modified ribonucleotides. Rna 19: 1684–1692

79. Saito K, Sakaguchi Y, Suzuki T, Suzuki T, Siomi H, Siomi MC (2007) Pimet, the Drosophila homolog of HEN1, mediates 2’-O-methylation of Piwi-interacting RNAs at their 3’ ends. Genes Dev 21: 1603–1608

80. Schmitt BM, Rudolph KL, Karagianni P, Fonseca NA, White RJ, Talianidis I, Odom DT, Marioni JC, Kutter C (2014) High-resolution mapping of transcriptional dynamics across tissue development reveals a stable mRNA-tRNA interface. Genome Res 24: 1797–1807

81. Schorn AJ, Gutbrod MJ, LeBlanc C, Martienssen R (2017) LTR-Retrotransposon Control by tRNA-Derived Small RNAs. Cell 170: 61–71.e11

82. Schorn AJ, Martienssen R (2018) Tie-Break: Host and Retrotransposons Play tRNA. Trends Cell Biol 28: 793–806

83. Sim G, Kehling AC, Park MS, Divoky C, Zhang H, Malhotra N, Secor J, Nakanishi K (2023) Determining the defining lengths between mature microRNAs/small interfering RNAs and tinyRNAs. Sci Rep 13: 19761

84. Singh U, Wurtele ES (2021) orfipy: a fast and flexible tool for extracting ORFs. Bioinformatics 37: 3019–3020

85. Su Z, Monshaugen I, Wilson B, Wang F, Klungland A, Ougland R, Dutta A (2022) TRMT6/61A-dependent base methylation of tRNA-derived fragments regulates gene-silencing activity and the unfolded protein response in bladder cancer. Nat Commun 13: 2165

86. Svendsen JM, Reed KJ, Vijayasarathy T, Montgomery BE, Tucci RM, Brown KC, Marks TN, Nguyen DAH, Phillips CM, Montgomery TA (2019) henn-1/HEN1 Promotes Germline Immortality in Caenorhabditis elegans. Cell Rep 29: 3187–3199.e3184

87. Tam OH, Aravin AA, Stein P, Girard A, Murchison EP, Cheloufi S, Hodges E, Anger M, Sachidanandam R, Schultz RM et al (2008) Pseudogene-derived small interfering RNAs regulate gene expression in mouse oocytes. Nature 453: 534–538

88. Tanaka S, Kunath T, Hadjantonakis AK, Nagy A, Rossant J (1998) Promotion of trophoblast stem cell proliferation by FGF4. Science 282: 2072–2075

89. Tareen A, Kooshkbaghi M, Posfai A, Ireland WT, McCandlish DM, Kinney JB (2022) MAVE-NN: learning genotype-phenotype maps from multiplex assays of variant effect. Genome Biol 23: 98

90. Tsherniak A, Vazquez F, Montgomery PG, Weir BA, Kryukov G, Cowley GS, Gill S, Harrington WF, Pantel S, Krill-Burger JM et al (2017) Defining a Cancer Dependency Map. Cell 170: 564–576.e516

91. Turelli P, Playfoot C, Grun D, Raclot C, Pontis J, Coudray A, Thorball C, Duc J, Pankevich EV, Deplancke B et al (2020) Primate-restricted KRAB zinc finger proteins and target retrotransposons control gene expression in human neurons. Sci Adv 6: eaba3200

92. Vainberg Slutskin I, Weingarten-Gabbay S, Nir R, Weinberger A, Segal E (2018) Unraveling the determinants of microRNA mediated regulation using a massively parallel reporter assay. Nat Commun 9: 529

93. Virtanen P, Gommers R, Oliphant TE, Haberland M, Reddy T, Cournapeau D, Burovski E, Peterson P, Weckesser W, Bright J et al (2020) SciPy 1.0: fundamental algorithms for scientific computing in Python. Nat Methods 17: 261–272

94. Volkman HE, Stetson DB (2014) The enemy within: endogenous retroelements and autoimmune disease. Nat Immunol 15: 415–422

95. Wang X, Ramat A, Simonelig M, Liu MF (2023) Emerging roles and functional mechanisms of PIWI-interacting RNAs. Nat Rev Mol Cell Biol 24: 123–141

96. Wang Z, Wang W, Cui YC, Pan Q, Zhu W, Gendron P, Guo F, Cen S, Witcher M, Liang C (2018) HIV-1 Employs Multiple Mechanisms To Resist Cas9/Single Guide RNA Targeting the Viral Primer Binding Site. J Virol 92

97. Watanabe T, Totoki Y, Toyoda A, Kaneda M, Kuramochi-Miyagawa S, Obata Y, Chiba H, Kohara Y, Kono T, Nakano T et al (2008) Endogenous siRNAs from naturally formed dsRNAs regulate transcripts in mouse oocytes. Nature 453: 539–543

98. Wynant N, Santos D, Vanden Broeck J (2017) The evolution of animal Argonautes: evidence for the absence of antiviral AGO Argonautes in vertebrates. Sci Rep 7: 9230

99. Xu H, Boeke JD (1990) Host genes that influence transposition in yeast: the abundance of a rare tRNA regulates Ty1 transposition frequency. Proc Natl Acad Sci U S A 87: 8360–8364

100. Yang A, Bofill-De Ros X, Shao TJ, Jiang M, Li K, Villanueva P, Dai L, Gu S (2019a) 3’ Uridylation Confers miRNAs with Non-canonical Target Repertoires. Mol Cell 75: 511–522.e514

101. Yang A, Bofill-De Ros X, Stanton R, Shao TJ, Villanueva P, Gu S (2022) TENT2, TUT4, and TUT7 selectively regulate miRNA sequence and abundance. Nat Commun 13: 5260

102. Yang Q, Li R, Lyu Q, Hou L, Liu Z, Sun Q, Liu M, Shi H, Xu B, Yin M et al (2019b) Single-cell CAS-seq reveals a class of short PIWI-interacting RNAs in human oocytes. Nat Commun 10: 3389

103. Yeung ML, Bennasser Y, Watashi K, Le SY, Houzet L, Jeang KT (2009) Pyrosequencing of small non-coding RNAs in HIV-1 infected cells: evidence for the processing of a viral-cellular double-stranded RNA hybrid. Nucleic Acids Res 37: 6575–6586

104. Yu S, Kim VN (2020) A tale of non-canonical tails: gene regulation by post-transcriptional RNA tailing. Nat Rev Mol Cell Biol 21: 542–556

105. Zhao Y, Yu Y, Zhai J, Ramachandran V, Dinh TT, Meyers BC, Mo B, Chen X (2012) The Arabidopsis nucleotidyl transferase HESO1 uridylates unmethylated small RNAs to trigger their degradation. Curr Biol 22: 689–694

106. Zheng G, Qin Y, Clark WC, Dai Q, Yi C, He C, Lambowitz AM, Pan T (2015) Efficient and quantitative high-throughput tRNA sequencing. Nat Methods 12: 835–837

107. Zhou H, Kimsey IJ, Nikolova EN, Sathyamoorthy B, Grazioli G, McSally J, Bai T, Wunderlich CH, Kreutz C, Andricioaei I et al (2016) m(1)A and m(1)G disrupt A-RNA structure through the intrinsic instability of Hoogsteen base pairs. Nat Struct Mol Biol 23: 803–810

